# Connectomes across temporal scales with simultaneous wide-field optical imaging and resting-state functional MRI

**DOI:** 10.64898/2026.02.01.703149

**Authors:** Wen-Ju Pan, Lauren Daley, Lisa Meyer-Baese, Shella D. Keilholz

## Abstract

Resting-state functional MRI (rs-fMRI) is a cornerstone of human brain research, yet its interpretation is complicated by its sensitivity to the slow hemodynamic response that obscures the organization of neural activity across faster time scales. Here we use simultaneous wide-field optical imaging (WOI) and rs-fMRI to directly examine the relationship between neural and hemodynamic functional connectomes across time scales. We show that much of the large-scale spatial structure is preserved across modalities, across time scales, and across frequencies. Although rs-fMRI robustly captures time-averaged neural activity, time-resolved rs-fMRI estimates of functional connectivity exhibit significantly greater variability, which partially reflects sensitivity limitations. Hemodynamic WOI signals maintain greater similarity to neural activity than rs-fMRI, although their fidelity is reduced at high frequencies. Together, our findings demonstrate that the time-averaged spatial structure of neural activity is faithfully represented in hemodynamics and rs-fMRI; provide insight into the reliability of time-resolved rs-fMRI across temporal scales; and establish a multimodal framework for validating features of dynamic brain activity.

**Teaser:** Spatial patterns of neural activity present across time scales are largely preserved in hemodynamics measured with optical imaging and rs-fMRI.

## Introduction

Like other complex systems, the brain exhibits structured intrinsic activity at a wide range of spatial and temporal scales, from millisecond-long events at a single synapse to slow fluctuations of global activity. Resting-state functional magnetic resonance imaging (rs-fMRI) based on the blood oxygenation level-dependent (BOLD) signal (*1*) is the most widely-used tool for the investigation of the organization of intrinsic brain activity in humans, but because the BOLD signal arises from the relatively slow hemodynamic response to neural activity, it obtains a filtered, low-frequency representation of the neural signal. As a result, it remains unclear which aspects of large-scale coordinated activity are faithfully captured by rs-fMRI, particularly for increasingly popular time-resolved analysis methods (*2–4*).

Multimodal studies using electrophysiology and rs-fMRI have confirmed that the BOLD fluctuations reflect the underlying neural activity (*5–12*), although measurements were typically averaged over many trials and were made from only a few locations, mostly in sensory cortex. Wide-field optical imaging (WOI) fills this gap by enabling cortex-wide readout of neural signals via genetically encoded calcium indicators and, in parallel, detection of hemodynamic signals via reflectance imaging (*10, 13, 14*). Simultaneous acquisition of WOI and rs-fMRI is technically challenging but has been successfully implemented, making it possible to directly compare the spatial organization of neural activity and the BOLD response in a time-locked manner (*15, 16*). This advance enables us to address three fundamental questions for the first time. First, to what extent is large-scale spatial structure preserved across modalities, and does it manifest similarly in WOI and rs-fMRI readouts of hemodynamics? Second, is this large-scale spatial structure specific to particular frequencies of neural activity? Finally, to what extent and on what time scales does time-resolved analysis of rs-fMRI data capture true reconfigurations of neural activity? The answers to these questions are essential for interpreting time-averaged and time-resolved rs-fMRI in humans, where no direct measurements of neural activity can be made.

We address these foundational questions using a newly-developed, highly sensitive method for simultaneous WOI and rs-fMRI in mice expressing GCaMP6f in excitatory neurons (*16*). Fluorescence (neural) and reflectance (total hemoglobin, HBT) WOI are acquired concurrently at 50 Hz during acquisition of whole-brain rs-fMRI at 1 Hz, and functional connectivity (FC) matrices are computed at the group and individual levels for all modalities using a range of frequencies and window lengths. This multimodal, multiscale approach allows us to directly examine for the first time how well the spatial structure of neural activity is reproduced in WOI and rs-fMRI measurements of hemodynamics across temporal scales.

Our results demonstrate that much of the spatial structure of brain activity is preserved across modalities and scales. Core spatial patterns of cortical activity are remarkably consistent across time scales in the neural signal and largely preserved in WOI measurements of hemodynamics, reinforcing prior observations of similar motifs of activity across time scales (*17*). A notable breakdown in the ability of WOI hemodynamics to capture the spatial structure of neural activity occurs for the highest frequency band, consistent with the inherent lowpass filter of the vascular response. Another key finding is that rs-fMRI robustly captures the spatial structure of time-averaged FC but gradually loses fidelity to the neural signal as time windows become shorter. Together, these results provide new insight into the ability of rs-fMRI to reflect the spatial organization of neural activity, enable validation of dynamic analyses against a ground truth, and establish a multimodal framework for understanding functional connectomes across temporal scales.

## Results

### Simultaneously-acquired rs-fMRI and WOI maps of functional connectivity

High-quality data were consistently obtained for both WOI and rs-fMRI. Data quality is examined in more detail in (*16*), but an example of FC in the barrel network across modalities for a single scan from one mouse is shown in **Figure 1**. Even at the individual level, the detected network exhibits the bilateral structure typically observed in group level FC maps. Note that for Figure 1, unlike for the rest of the studies, FC was obtained on unparcellated data using ICA to better depict the detailed spatial correspondence across modalities.

**Figure 1.**
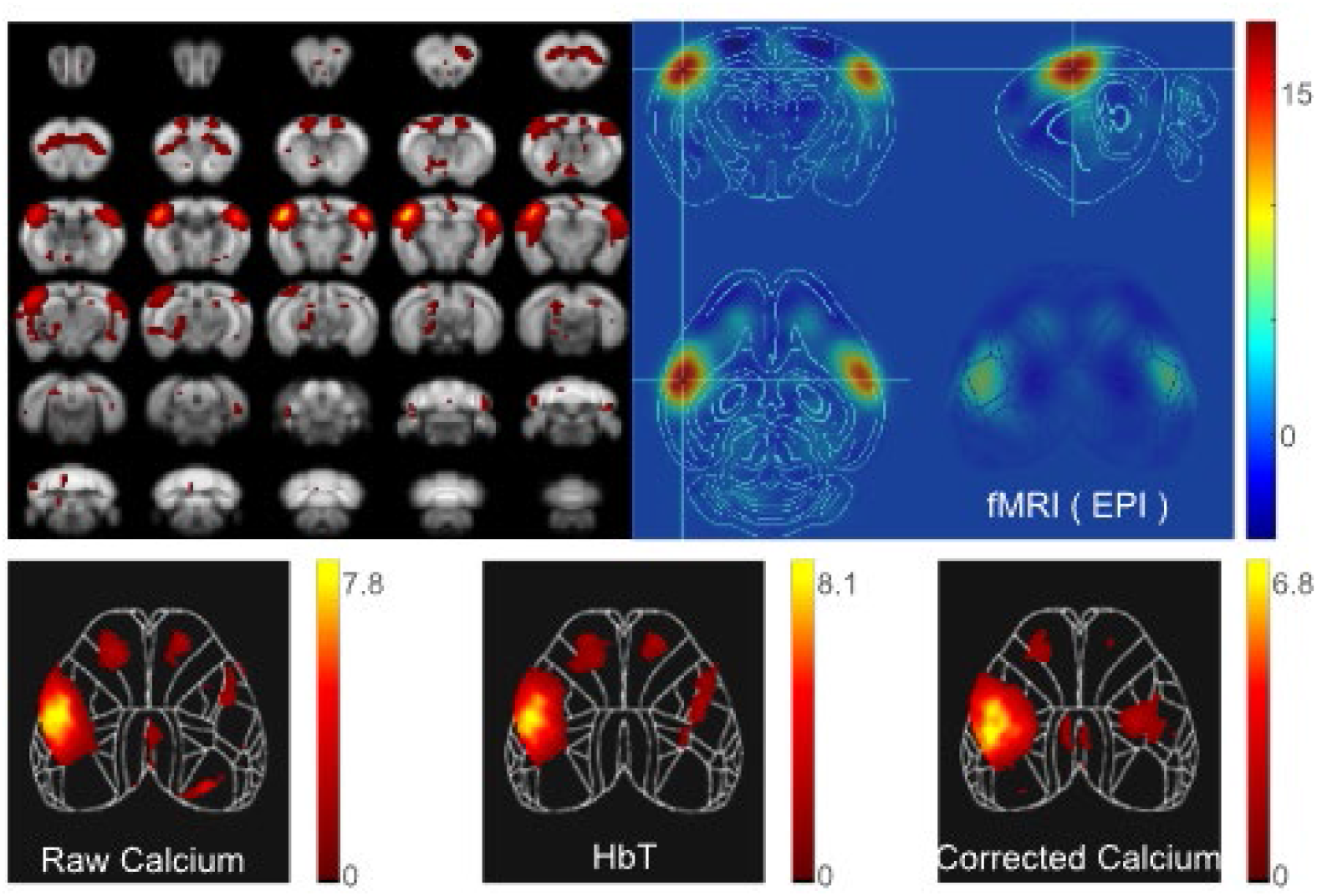
High-quality FC maps from all modalities. **Top row**) The barrel cortex network obtained using ICA on unparcellated rs-fMRI data from one scan in one mouse shown on anatomical MRI and the Allen atlas, including a cortical surface projection for comparison to WOI in the bottom right corner of the righthand panel. All values are shown as z scores. **Bottom row**) ICA-based FC from same scan in the unparcellated raw fluorescence signal, HBT, and fluorescence signal after correction for hemodynamic contamination. For all modalities, a bilateral sensory network is observed.

### Similarity of time-averaged FC across modalities

Cortical FC matrices were calculated for the parcellated rs-fMRI, fluorescence and HBT data for each mouse for each modality (**Figure 2**). We first consider the group average results. At the group level, the average fluorescence and HBT FC matrices are nearly identical to the eye (spatial correlation 0.97). The amplitudes of the correlations in the FC matrix are weaker in rs-fMRI, particularly for connections that have strong anticorrelation in the WOI FC matrices. However, rs-fMRI recovers the dominant bilateral network organization present in the WOI matrices (spatial correlation of 0.73 for rs-fMRI to fluorescence, 0.76 for rs-fMRI to HBT). Two notable aspects of these results are the strong similarity of the fluorescence and HBT FC matrices, which supports a consistent neurovascular coupling across the cortex; and the relative equivalence of fluorescence and HBT FC in terms of spatial correlation to the rs-fMRI FC. Because both HBT and rs-fMRI reflect hemodynamics rather than neural activity, FC from rs-fMRI might plausibly resemble HBT FC more closely than neural FC, but the current findings suggest that this is not the case and that both hemodynamic modalities closely reflect neural activity.

**Figure 2.**
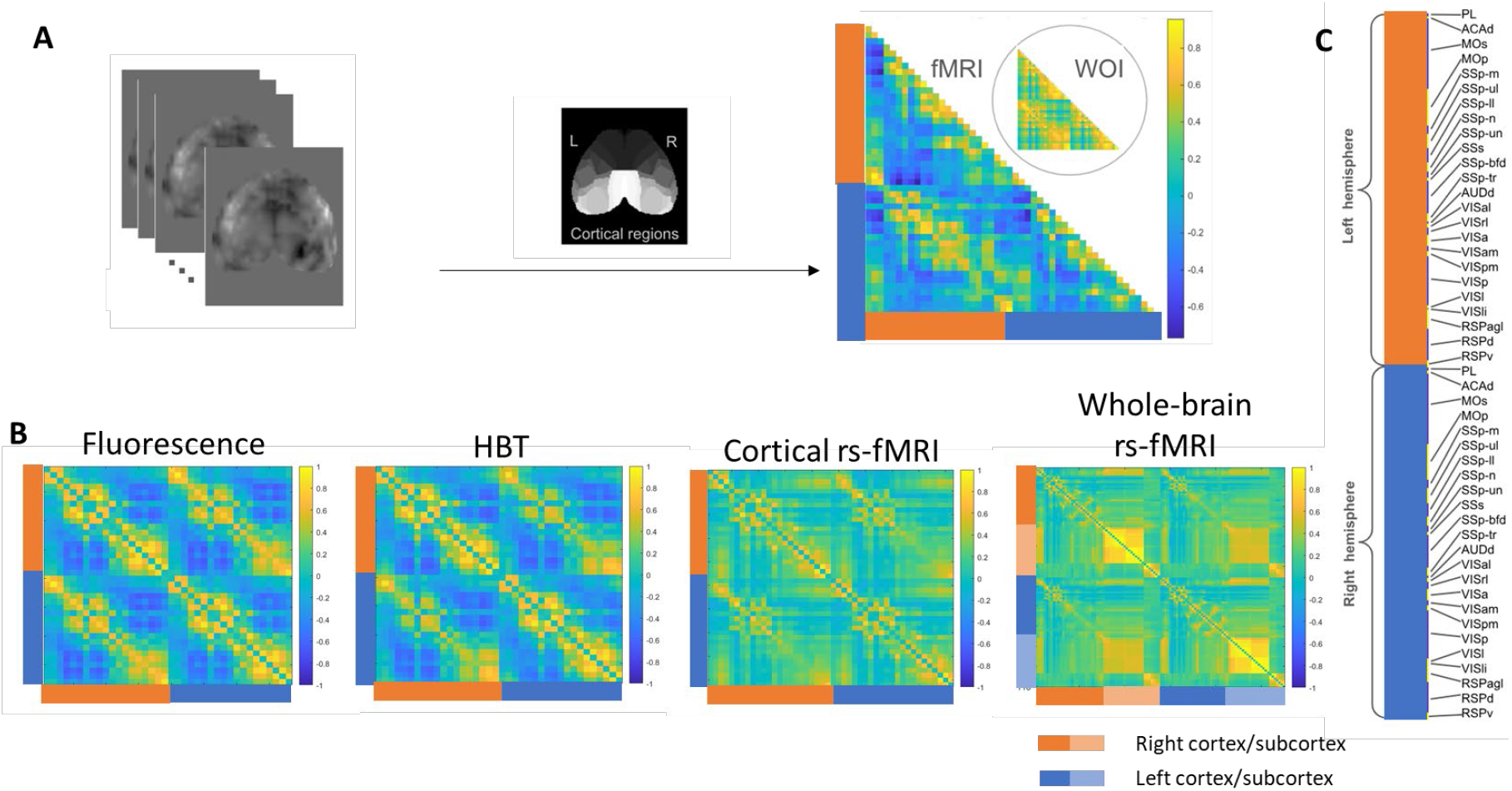
Group level FC matrices for all modalities. **A**) For rs-fMRI data, signal from the 3D volume corresponding to each cortical parcel is averaged to create the FC matrix. For WOI data, signal from the 2D area corresponding to each cortical parcel is used. **B**) Group average FC matrices for fluorescence, HBT, cortical rs-fMRI, and whole-brain rs-fMRI. Left and right hemispheres are indicated by orange and blue bars, respectively. FC matrices for fluorescence and HBT are highly similar. The FC matrix for cortical rs-fMRI generally contains weaker correlation, particularly in areas of strong anticorrelation in the WOI data. However, the spatial pattern of FC is largely conserved in rs-fMRI for both inter- and intrahemispheric connections. **C**) List of cortical parcels for each hemisphere.

The similarity of FC matrices obtained with different modalities was also examined at the individual level. Spatial correlation between FC matrices was calculated for each scan (**Figure 3A**). FC matrices from HBT and fluorescence maintained a high level of similarity even at the individual level, with an average spatial correlation of 0.81 (compared to 0.97 at the group level). Spatial correlation was significantly lower for rs-fMRI FC matrices with either WOI modality at the individual level (approximately 0.4-0.5, compared with 0.73-0.76 at the group level) but the strongest features (e.g., interhemispheric correlation) were typically preserved.

**Figure 3.**
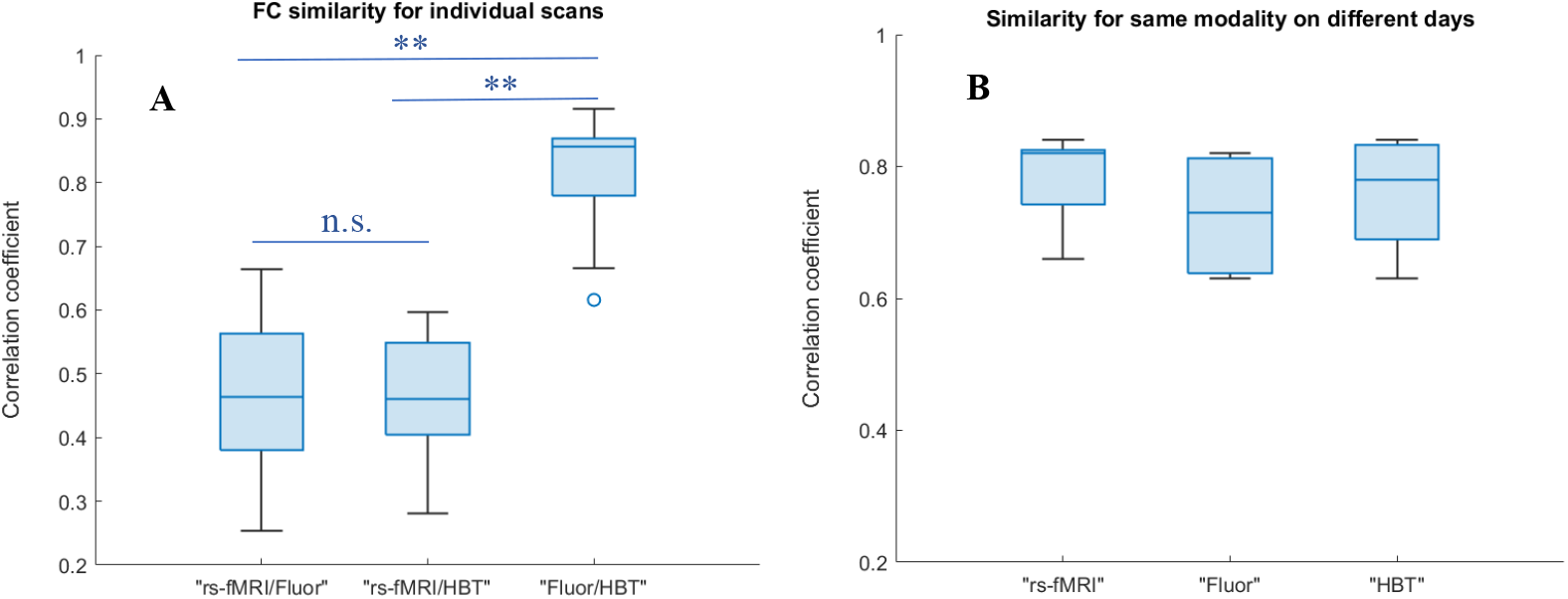
Spatial similarity of FC across modalities and across days. Similarity between time-averaged FC matrices from different modalities for the same scan (**A**) and between the same modalities on different days (**B**, no significant differences). ** p<0.001, Bonferroni correction.

Three mice were scanned repeatedly on different days, for a total of 7 scans. For these mice, we examined the similarity of the FC matrices obtained using the same modality during different scans. FC matrices for all modalities in this small group of animals exhibited comparable strong correlation across days (**Figure 3B**).

### Community structure of FC matrices across modalities

To provide a complementary assessment of spatial similarity across modalities, the Louvain community detection algorithm was applied to the group-level FC matrices to obtain an estimate of the modularity (q) and the community assignment for each parcel. Group level modularity was 0.26 for rs-fMRI of the cortex, 0.17 for rs-fMRI of the whole brain, 0.43 for WOI-fluorescence, and 0.41 for WOI-HBT, with the lower modularity for rs-fMRI possibly arising from the reduced amplitude of anticorrelation noted previously. Three large communities were obtained for fluorescence and HBT, two for cortical rs-fMRI, and three for whole-brain rs-fMRI. As shown in **Figure 4**, community assignments for cortical parcels were identical for fluorescence and HBT images. For cortical rs-fMRI, communities 1 and 3 for the WOI data were merged into a single cluster. For all modalities, clusters were bilaterally symmetric, indicative of the strong bilateral FC between homologous areas in the left and right hemispheres.

**Figure 4.**
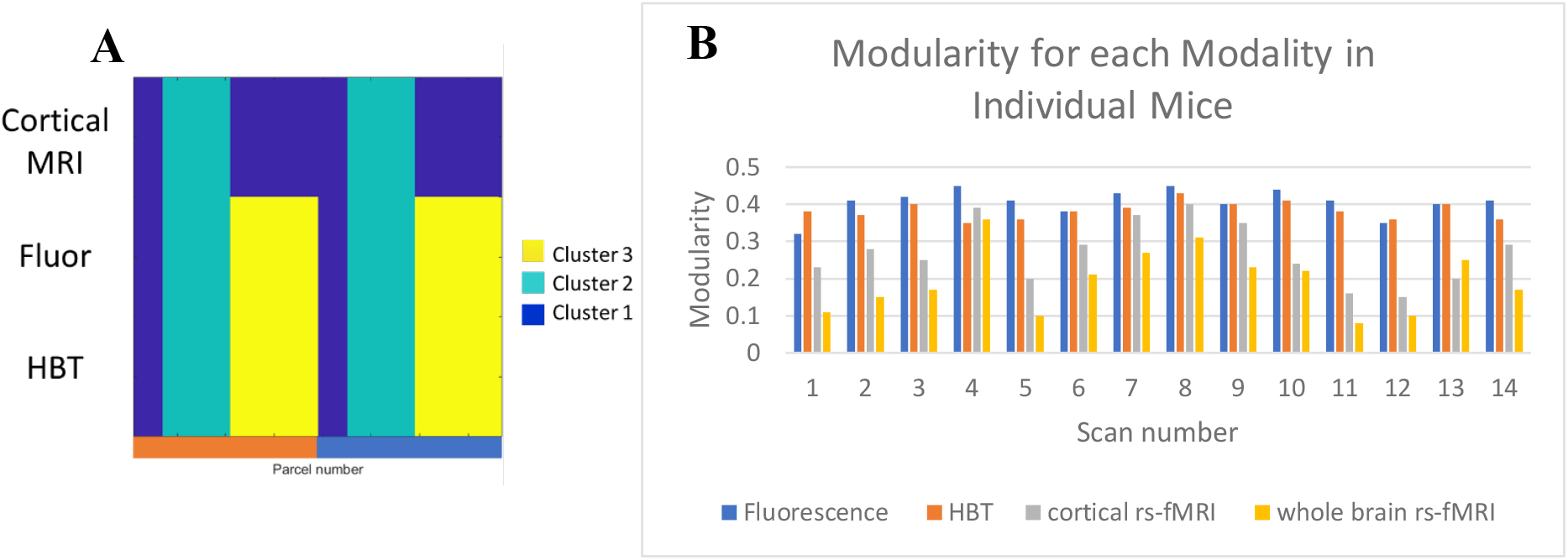
Community structure and modularity for all modalities. A) Community structure for cortical rs-fMRI, fluorescence, and HBT based on the group average FC matrices. For each modality, cortical parcels are arranged along the horizontal axis, and the color at each point indicates the community assignment for that parcel. Communities are identical for fluorescence and HBT, and the primary difference for rs-fMRI is that two clusters merge into one. B) Modularity calculated for each modality for FC matrices from each individual scan. WOI modularity is typically higher than cortical rs-fMRI modularity, and cortical rs-fMRI modularity is higher than whole-brain rs-fMRI modularity.

Modularity was also calculated for the FC matrices for the three modalities from each individual scan. As shown in **Figure 4**, the values for rs-fMRI were lower than the values for WOI-fluorescence and HBT, which were comparable to each other. The average modularity values calculated at the individual level were roughly equivalent to those calculated on the group matrix. For cortical rs-fMRI, modularity was 0.27 +/- 0.08; for whole-brain rs-fMRI, 0.2 +/-0.08; for fluorescence, 0.41+/-0.04; and for HBT, 0.38 +/- 0.02. Modularity for rs-fMRI was significantly lower than modularity for either WOI modality, but no significant difference was observed for fluorescence and HBT. The percent of brain parcels assigned to the same community in rs-fMRI and fluorescence averaged 65 +/- 12%; for rs-fMRI and HBT, 68 +/- 11%, and for fluorescence and HBT, 79 +/- 15%. These findings demonstrate that the increased modularity observed in WOI at the group level is maintained at the individual level and provide further evidence that important aspects of the spatial structure of the neural signal is preserved across modalities, even at the level of individual scans.

### Frequency dependence of spatial structure

Neural activity spans a wide range of frequencies, while the accompanying hemodynamic fluctuations are confined to a much smaller range. To examine the frequency dependence of the spatial structure captured by FC matrices, fluorescence and HBT images were filtered into infraslow (0.01-0.1 Hz), slow (0.1-1 Hz), and delta (1-4 Hz) bands, and FC matrices were created at the group and individual level for each band. Only the infraslow band was examined in rs-fMRI due to the limitations imposed by the 1 Hz sampling rate. However, prior studies have shown that coherence between electrophysiological recordings and rs-fMRI is mostly confined to the infraslow frequency range during isoflurane anesthesia (*7*).

At the group level, the spatial structure for fluorescence as captured by the FC matrix is highly similar across all frequency bands (**Figure 5**; spatial correlation of 0.96 for infraslow and slow bands; 0.93 for infraslow and delta). These findings are consistent with prior work that found similar spatial motifs occurring at different time scales (*17*) and demonstrate that the large-scale spatial structure of neural activity is preserved across the frequencies observable with calcium indicators.

**Figure 5.**
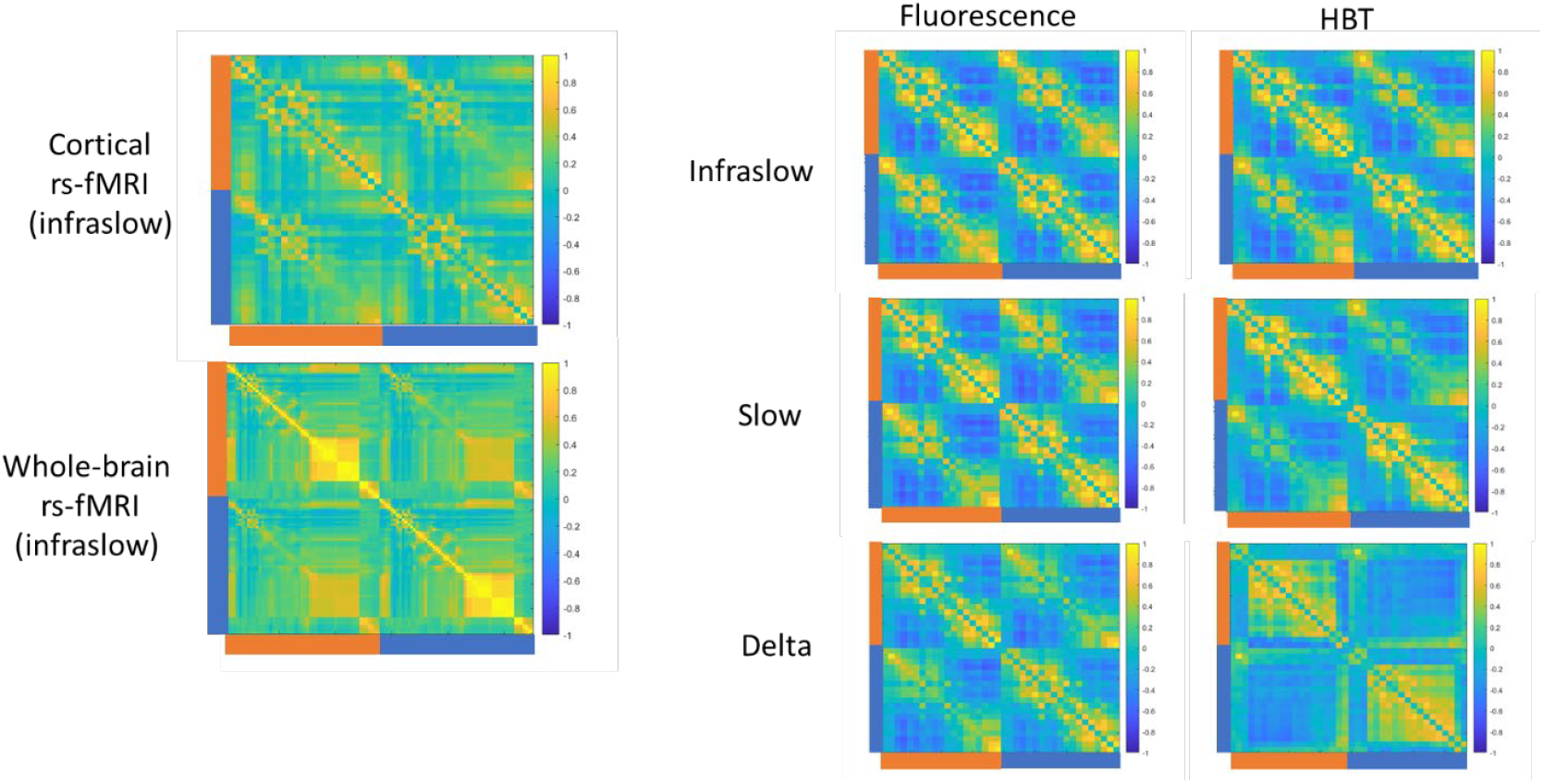
Group level FC for different frequency bands of WOI. Group level FC matrices for fluorescence and HBT filtered into infraslow, slow, or delta frequency bands. FC is similar across bands for fluorescence but differs for the delta band in HBT, with a prominent loss of interhemispheric connectivity. Group level cortical and whole-brain FC matrices from the infraslow band are shown at left for comparison.

For HBT, unlike fluorescence, the delta band exhibits a loss of structure, specifically in interhemispheric FC (spatial correlation of 0.97 for infraslow and slow bands; 0.52 for infraslow and delta bands). This is not surprising given the inherent low pass filter of the hemodynamic response. Modularity, however, is similar for the three frequency bands for both fluorescence and HBT: 0.43 and 0.41 for the infraslow band, 0.45 and 0.39 for the slow band, and 0.45 and 0.45 for the delta band. The similarity of the modularity for HBT across frequency bands despite the clear differences in the spatial structure of the delta band suggests that modularity, while providing a useful summary of overall structure, may not be a sensitive measure for some aspects of spatial reorganization. Given the lack of change in neural FC, the notable loss of interhemispheric connectivity in the HBT FC matrix should be interpreted as the effect of the known low-frequency filtering inherent in the hemodynamic response rather than an alteration of spatial structure across frequency scales.

### Preservation of spatial structure in hemodynamics for time-resolved analysis

In addition to encompassing a wide range of frequency bands, neural activity also demonstrates time-varying spatial patterns that average over the course of data acquisition to create the spatial structure of FC measured in a typical rs-fMRI scan, which we will refer to as time-averaged FC. Many time-resolved analysis methods have been proposed to characterize the time-varying patterns that are combined to create time-averaged FC, the simplest of which is windowed FC. In this approach, image time courses are divided into shorter windows prior to the calculation of the FC matrices, so that instead of a single FC matrix per scan, each scan provides a time series of FC matrices. To examine the ability of hemodynamic modalities to capture changes in the spatial structure of neural activity over time, FC matrices were calculated for infraslow band for rs-fMRI, HBT, and fluorescence signals using windows of 15, 30, or 60 seconds. Substantial variability in FC across windows was evident for all modalities. As an example, FC matrices from a randomly-chosen set of three consecutive 30 second windows are shown from one scan in **Figure 6A**. Differences in the spatial structure for all three modalities were observed over time, with the most prominent changes in the rs-fMRI FC matrices. To provide insight into the relative stability of each connection, the standard deviation for each connection in the FC matrix was calculated across all windows from all animals for each modality (**Figure 6B**). A core group of intra- and inter-hemispheric connections exhibits relatively low variability over time, evidence of a preserved backbone of spatial structure across modalities and time scales.

**Figure 6.**
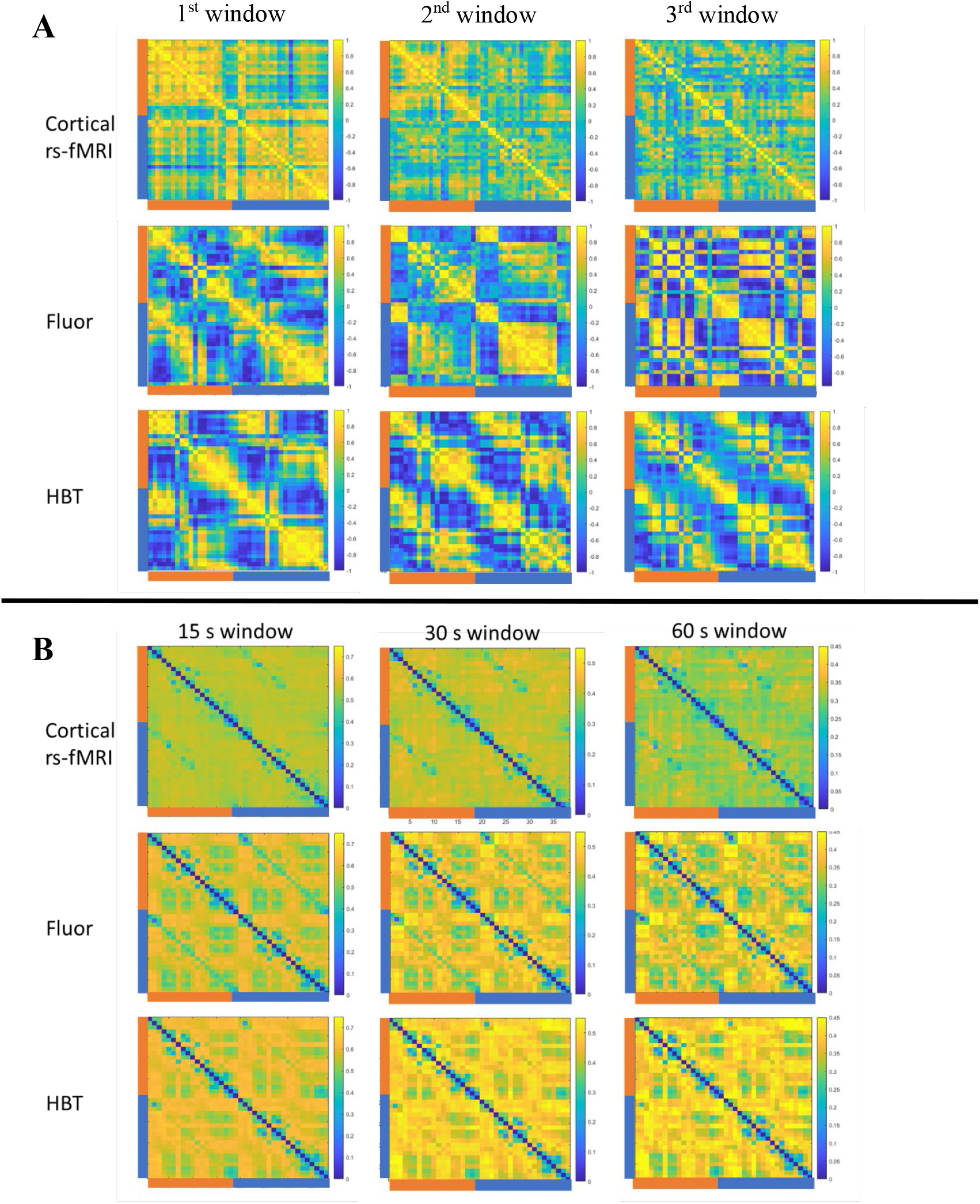
Time-varying FC across modalities. **A**) FC matrices created from a randomly-selected set of three consecutive 30 second windows from one scan. The rs-fMRI matrices exhibit the greatest spatial variability but changes are also evident in both WOI matrices. **B**) The standard deviation for each connection across all windows and all animals for infraslow band activity from each modality. A subset of connections exhibit relatively low variability over time for all modalities. Note that scales are different for each window length (but consistent across modalities) to allow visualization of structure.

To characterize the variability present in the windowed FC matrices within each modality, spatial correlation was calculated pairwise for all windows (**Figure 7A**). Variability across windows is challenging to interpret, as both true neural reconfiguration and fluctuations arising from noise contribute to the variability observed. However, variability in the fluorescence FC might be considered an upper bound for true neural variability. In concordance with this framing, similarity across windows was highest for fluorescence for any given window length, although it was only significantly greater than the similarity across windows for HBT for the 60 second window in the infraslow band. Despite the small but consistent difference, the similarity across windows for HBT and fluorescence was roughly comparable. For rs-fMRI, however, the similarity was significantly lower for all window lengths, suggesting an additional source of variability.

**Figure 7.**
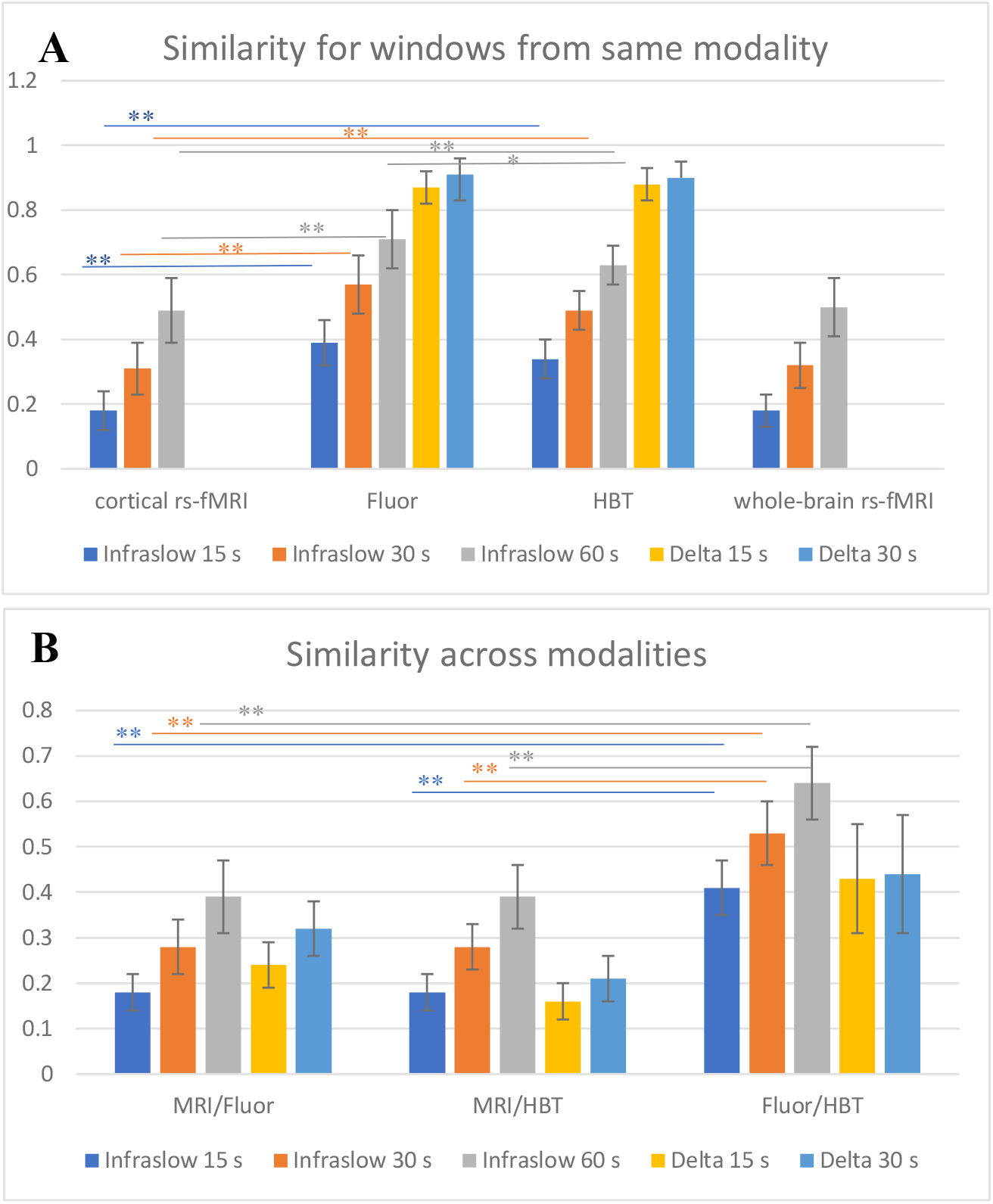
Spatial similarity for windows within and across modalities. **A**) Spatial similarity measured as pairwise spatial correlation of FC matrices for each modality for multiple window lengths and frequency bands. For all modalities, similarity increases as window length increases, most notably for the infraslow band. **B**) Spatial similarity across modalities for the same window. Similarity increases with window length for both frequency bands for rs-fMRI compared to either WOI modality. Similarity is highest for fluorescence and HBT, and comparable for rs-fMRI with either WOI modality. * p<0.05; **p<0.001; Bonferroni correction

Similarity across windows increased as window length increased for all modalities, which should be expected as longer windows more closely approximate the time-averaged spatial structure and tend to average out the effects of genuine spatial reconfigurations along with transient noise. To determine whether the similarity across windows was dependent upon the frequencies used for WOI, we examined FC matrices created from the highest frequencies (delta band) using windows of 15 and 30 seconds. The delta band FC matrices exhibited higher similarity over time than infraslow FC and less dependence upon window length. The similarity across windows in the delta band was roughly comparable to the similarity of scans obtained from the same animals on different days. Together with the nearly equivalent results for the 15 and 30 second windows, this suggests that these short windows of delta band activity already approach the steady state spatial structure observed with time-averaged measurements (in concordance with a prior optical study (*14*)). This motivated us to examine extremely short windows for the WOI data, to determine whether faster variability could be detected. Using a window length of 1 second (containing 1-4 cycles of activity in the delta band; comparable to a single rs-fMRI time point), we calculated the FC matrices for fluorescence and HBT images. The similarity of the FC matrices obtained across 1 second windows for fluorescence was lower than the similarity obtained from longer windows of delta band activity (0.44±0.07), suggesting that these shorter windows may begin to resolve changes in spatial patterns of neural activity at these higher frequencies, although variability due to noise also increases due to the reduced number of data points used to calculate FC. Similarity across windows was higher but more variable for the HBT data (0.57±0.15), which likely reflects the loss of information observed at delta band frequencies and a resulting smoothing of patterns across time.

### Spatial patterns of neural activity at different time scales

To better understand whether the same spatial patterns of neural activity arise at different temporal scales, we compared the FC matrices created from infraslow fluorescence with 30 s windows to FC matrices created from delta band fluorescence with 1 s windows. To account for the vastly different number of windows, k-means clustering (k=3) was applied to each modality and the average patterns for the resulting clusters were compared. As shown in **Figure 8**, for at least the two largest clusters, the similarity across temporal scales is strong, supporting prior reports of similar spatial patterns at different temporal scales.

**Figure 8.**
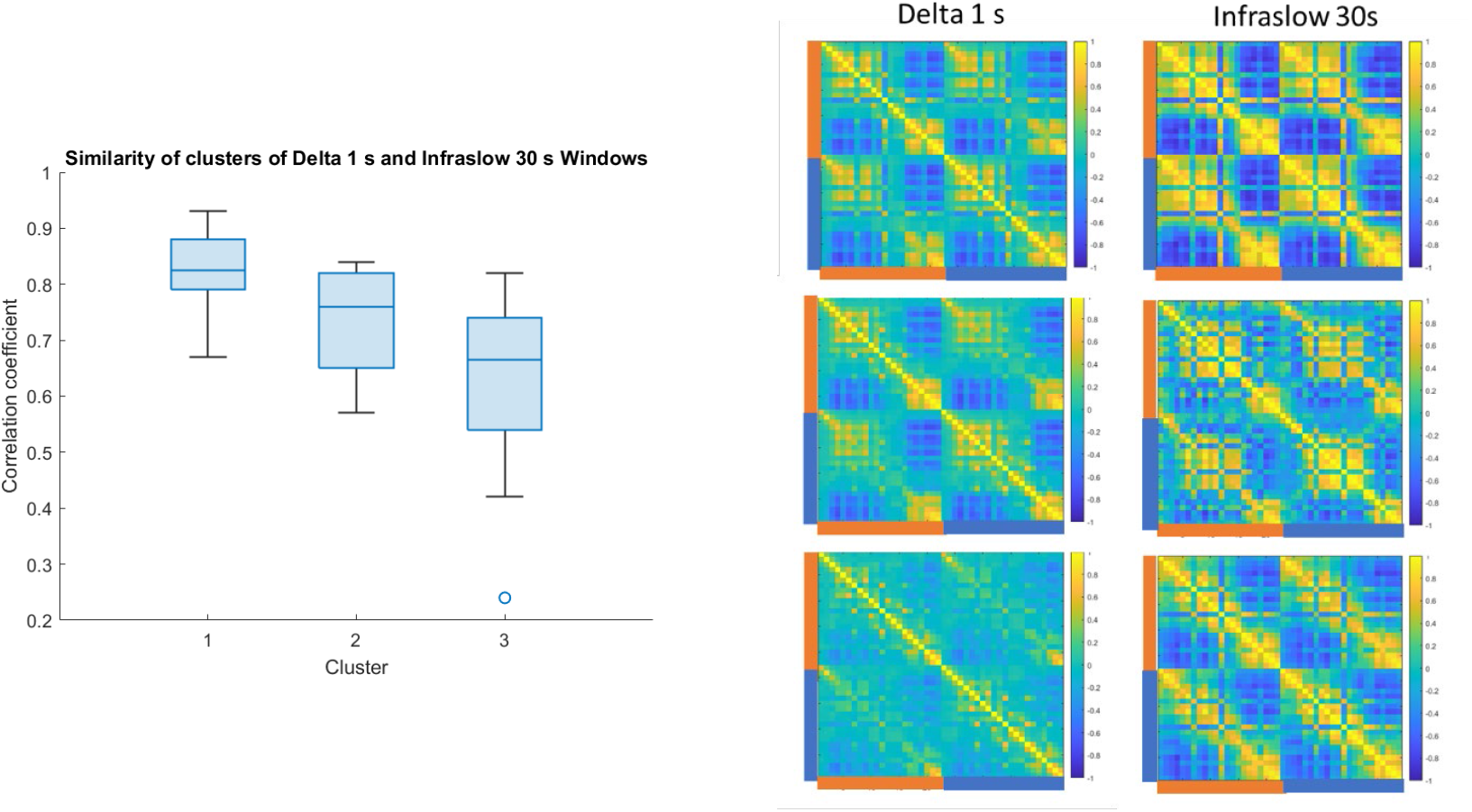
Spatial similarity of neural activity across window lengths. Similarity of the average FC pattern obtained by clustering FC matrices created from 1 second windows of delta band neural activity and 30 s windows of infraslow neural activity. The high spatial correlation indicates that similar patterns of activity occur at very different time scales. Mean images from each cluster for delta and infraslow activity are shown at right for one randomly-chosen mouse. Correlation values are typically weaker for delta band, particularly for the third cluster, but much of the spatial pattern is preserved.

### Cross-modality correspondence of spatial structure for different time scales

While the similarity of FC matrices for fluorescence across windows puts an upper bound on the variability due to changes in neural activity, it is also important to determine how well these changes are reflected in the accompanying hemodynamics. To determine how well HBT and rs-fMRI captured time-varying changes in neural FC, we examined the relationship between FC matrices obtained during the same time window with different modalities (**Figure 7B**). The FC matrices obtained from infraslow rs-fMRI were compared to WOI FC matrices obtained from either infraslow or delta bands for the same window. Spatial correlation for all frequency bands and window lengths was highest for fluorescence and HBT, with similarity increasing as a function of window length for the infraslow band. Similarity for either window length in the delta band was roughly comparable to that of the shortest window length in the infraslow band, and exhibits more variability than for other bands. This again is likely to reflect the loss of information in the hemodynamic signal from these bands.

Similarity for rs-fMRI and either WOI modality is markedly lower than for fluorescence and HBT, roughly half the HBT/fluorescence spatial correlation at any given window length. This suggests that much of the variability in the rs-fMRI signal arises not from its reliance on neurovascular coupling (which it shares with the HBT signal) but on additional sources of non-neural variability or its low sampling rate. To address the role of the low sampling rate in the introduction of variability, WOI data were downsampled from 50 Hz to 1 Hz after preprocessing but prior to the creation of FC matrices. Downsampling had no significant effect on the similarity across windows within a modality or the cross-modality similarity for each window (shown for 30 second windows of infraslow activity in **Supplemental Figure 2**), suggesting the low sampling rate of rs-fMRI alone is not sufficient to account for the variability observed.

To summarize the large-scale changes in spatial structure across windows, modularity was calculated for each windowed FC matrix from each modality using the range of window lengths and frequency bands described above. However, the average modularity changed little across conditions, and the correlation between time courses of modularity from different modalities was near zero. This again suggests that modularity is not ideal for capturing some of the changes observed in the spatial structure of the brain’s intrinsic activity.

To better understand how well time-resolved analysis of rs-fMRI captures the spatial structure of neural activity, we further investigated the relationship between windowed rs-fMRI and fluorescence FC matrices. For every rs-fMRI scan, there are windows with relatively high similarity to fluorescence FC from the same window, and other windows with relatively low cross-modality similarity. We examined the spatial patterns for each modality in windows with high (top 10%) and low (lowest 10%) spatial correlation. An example is shown in **Figure 9** for one mouse using the infraslow bands for both modalities and a 30 second window. In windows with high correlation, rs-fMRI and fluorescence both exhibit graded spatial structure that is loosely organized into areas of positive and negative correlation. At time points of low correlation, however, the rs-fMRI signal exhibits little structure, and the fluorescence signal is strongly divided into positive and negative clusters. The loss of structure in the FC matrix for rs-fMRI can be characterized using the distribution of correlation values within each FC matrix (see **Supplemental Figure 3**). While the example shown is from a single mouse, this pattern was observed consistently across mice. FC matrices created for the infraslow band windows with high and low correlation for all mice can be found in **Supplemental Figure 4**.

**Figure 9.**
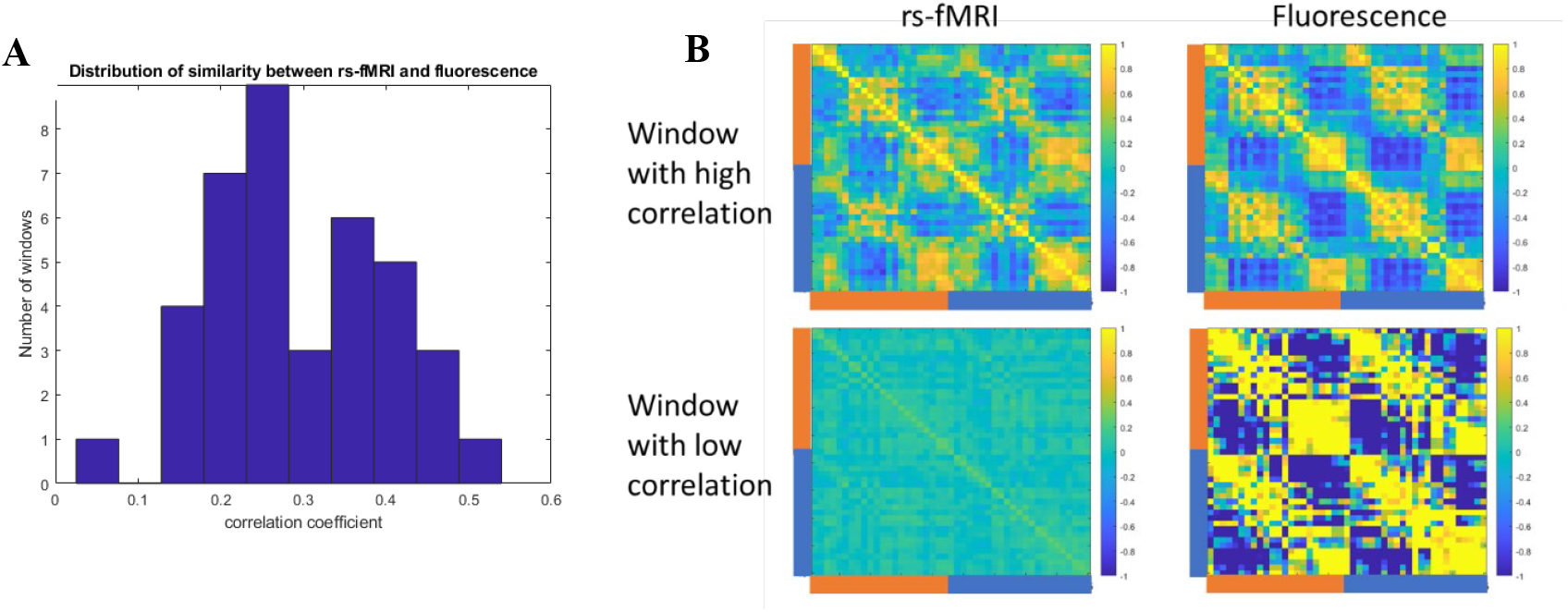
Time windows with high and low correlation between rs-fMRI and neural activity. A) The distribution of spatial correlation between FC matrices obtained from rs-fMRI and fluorescence in one mouse (infraslow band, 30 second window). Windows with particularly high or low similarity lie at the extremes of the distribution and were used to create the FC matrices in panel B. B) The average FC matrices obtained from the windows with high correlation and low correlation for rs-fMRI and fluorescence for the same scan. Notably, in windows with high correlation, rs-fMRI and fluorescence both exhibit graded spatial structure that is loosely organized into areas of positive and negative correlation. At time windows with low correlation, however, the rs-fMRI signal exhibits little structure, and the fluorescence signal is strongly divided into positive and negative clusters.

## Discussion

By combining WOI calcium and hemodynamic imaging with simultaneous rs-fMRI measurements, we were able to directly examine the relationship between neural activity and hemodynamics at different temporal scales. Notably, we found that the spatial structure of neural activity in the cortex is present across a range of temporal scales. This scale invariance may explain why the same structure is largely preserved in HBT in the infraslow and slow bands. The spatial patterns are diminished in the delta band in HBT due to the inherent lowpass vascular filter. Time-averaged FC from rs-fMRI robustly captures neural FC, but the correspondence to neural FC is weaker for windowed approaches. Together, these findings provide insight into how well rs-fMRI can represent spatial patterns of neural activity, point toward improved interpretations of FC based on rs-fMRI, and establish a multimodal framework for the validation of time-resolved rs-fMRI analysis.

### Hemodynamic modalities reflect the spatial structure of neural activity across the cortex

Spatial structure was highly similar for fluorescence and HBT at both the group and individual level, evidence of consistent neurovascular coupling across the cortical areas examined. It should be noted that the raw fluorescence signal is contaminated by absorption related to hemodynamics (*18*), but regression-based correction of concurrently-acquired reflectance minimizes the hemodynamic component, making it unlikely that shared hemodynamic fluctuations (rather than neurovascular coupling) account for the similarity of the HBT and fluorescence signals (*16*). The attribution of similarity to neurovascular coupling is further supported by the fact that the FC matrix of fluorescence in the delta band is highly similar to those obtained from the slow and infraslow bands, while the HBT signal loses structure at delta band frequencies. Much of the spatial structure of the neural activity was preserved in the rs-fMRI, although the rs-fMRI FC matrices were less similar to neural FC than the HBT matrices were. The robust similarity of FC from fluorescence and HBT demonstrates that neurovascular coupling is not the sensitivity-limiting factor for rs-fMRI studies, and puts an optimistic upper bound on the fidelity to neural activity that might eventually be achieved with rs-fMRI.

### Reduced similarity of rs-fMRI FC to neural FC in windowed analysis

Shorter time windows led to higher variability in the spatial structure obtained from all modalities. This has been shown for rs-fMRI before (*19–21*), but interpretation was challenging because the increased variability could arise from greater sensitivity to short-lived neural activity instead of or in combination with increased variability from shorter time samples. As is clear from Figure 7, variability in rs-fMRI has a substantial component that is not explained by variability in neural activity. Even if one assumes that all of the variability in fluorescence-based FC arises from time-varying spatial patterns rather than noise, the similarity across time windows for rs-fMRI is roughly half to two thirds of the spatial correlation for windows of fluorescence FC. Nor can the difference be explained by hemodynamic coupling, as the spatial similarity for HBT is only slightly lower than for fluorescence. Instead, we interpret these findings as showing that neurovascular coupling is consistent but that rs-fMRI is operating near a critical sensitivity level. This is further supported by the observation that the time windows with lowest cross-modality similarity between rs-fMRI and WOI all lack discernible spatial structure in the rs-fMRI FC matrices, suggesting that the spatial structure of the neural activity is somehow buried by the occurrence of unstructured noise.

Note that no hemodynamic delay was incorporated when comparing the two hemodynamic modalities to the fluorescence measures of neural activity. A delay of ∼2 s is unlikely to have much influence on the characterization of infraslow activity over 30 s or more, but could have caused a slight underestimation of the correlation between modalities in the short 15 second windows. However, the similarity of FC for HBT and fluorescence across all parameters examined suggests the effects are minimal.

### Implications for interpretation of rs-fMRI

One of the key but inadequately tested assumptions underlying rs-fMRI is that neurovascular coupling provides a consistent representation of the underlying neural activity. Neurovascular coupling has been reliably observed in sensory areas where most studies that link neural activity and hemodynamics are performed, but prior studies have shown that coupling may be different in other regions of the brain under certain conditions (*22–24*). The similarity observed across infraslow fluorescence and HBT FC matrices throughout this study demonstrates that neurovascular coupling across the entire cortex is adequate to faithfully represent the large-scale spatial structure of neural activity, even when windowed analysis is applied. The FC matrices from rs-fMRI preserve much of this structure for time-averaged analysis but become progressively less similar to neural FC as window length decreases. Because the BOLD signal is closely related to cerebral blood volume, the source of HBT contrast, the difference in sensitivity is likely to arise from the sparser sampling of the rs-fMRI data and/or additional sources of noise that reduce the quality of FC estimates. HBT should have improved sensitivity for FC relative to rs-fMRI due to its smaller pixel size and faster sampling rate, which essentially means that noise in the time course from each brain parcel is reduced by substantial averaging. Moreover, the BOLD signal is known to be contaminated with noise from motion and physiological processes (*25–28*). While these noise sources were minimized in the anesthetized mice imaged for this experiment, residual contributions could remain. Our work shows that despite the reduced sensitivity, rs-fMRI captures the large-scale spatial structure of neural activity quite well for time-averaged studies, but less well when shorter time windows are used.

The observation that poor correspondence to neural FC typically manifests as a lack of structure in the rs-fMRI FC matrix, along with the consistent similarity observed between HBT and fluorescence FC, is an encouraging indication that rs-fMRI studies could be optimized so that neural FC is more reliably represented in short time windows. For example, high field scanners and cryogenic RF probes boost SNR and could enhance the sensitivity of rs-fMRI. Postprocessing approaches such as NORDIC (*29, 30*) may also improve sensitivity. The advent of simultaneous WOI/rs-fMRI can provide a ground truth against which these optimizations can be tested.

With interest increasing in time-resolved analysis methods that capture the evolution of whole-brain activity, it is important to extend this validation to the dynamic evolution of the entire connectome. A number of time-resolved methods for the analysis of rs-fMRI data have been developed (*2–4*). Windowed correlation was one of the earliest time-resolved approaches (*19, 31, 32*), and its many drawbacks are well-documented (*20, 21, 33*). Nevertheless, prior work has shown that some information about coordinated neural activity is preserved in sliding window correlation for rs-fMRI (*34, 35*). The question of how much information about the spatial structure of neural activity is preserved in windowed FC matrices created from rs-fMRI previously lacked an answer because no method could obtain simultaneous measurements of neural activity with sufficient spatial coverage and resolution. The development of rs-fMRI/WOI made this experiment possible. As suspected, longer windows provide better estimates of the underlying neural activity. Moreover, it may be possible to identify windows with high vs. low information about neural activity based upon properties of the signal or its spatial distribution.

The very preliminary comparison of windows with high and low similarity to neural FC suggests that a loss of spatial structure in the FC matrix is rarely connected to a similar loss of neural FC and is likely instead to indicate obscuration with noise. This is similar to the observation behind another common time-resolved analysis, coactivation patterns (CAPs), that FC is driven primarily by a few high-amplitude events (*36–38*). Moreover, some connections are detected more reliably than others, which may provide another avenue for estimating fidelity of rs-fMRI FC when ground truth is unknown. Further work is needed, but the current results provide reason for cautious optimism that some aspects of time-varying neural FC can be extracted with rs-fMRI.

### Frequency dependence and multiscale motifs

An important finding from this study was the scale-invariance of the spatial structure of cortical activity. Very little difference was observed for fluorescence FC created from different frequency bands, suggesting that the same spatial structure at higher frequencies is nested within similar spatial patterns at lower frequencies. An intriguing prior WOI study showed that dynamic patterns of neural activity at higher frequencies (3-6 Hz) were mostly recapitulated in the infraslow (<1 Hz) range, more evidence that similar spatial patterns can occur at multiple temporal scales (*17*). This type of scale invariance might explain why rs-fMRI can capture much of the structure of neural activity, despite its inherently slow time scale.

The infraslow frequencies are of special interest given that this band gives rise to the strongest coherence between the BOLD signal and electrophysiological measurements of neural activity. Infraslow electrical activity itself is poorly understood, and previous measurements were limited to a few sites (*7, 12, 39, 40*). The simultaneous WOI/rs-fMRI study reported here extends this characterization across the cortex, and shows that FC measured with rs-fMRI is closely linked to the underlying infraslow neural FC. Moreover, neural FC exhibits the same spatial structure across infraslow, slow, and delta bands. Higher frequency bands were not examined due to limitations of the fluorescence indicator but future work with faster calcium or voltage indicators would enable the examination of FC above the delta band.

### Relevance of mouse FC to human FC

WOI can image neural activity and hemodynamics concurrently across most of the cortex with much higher spatial and temporal resolution than rs-fMRI, enabling unprecedented insight into how well spatial patterns of neural activity are captured with rs-fMRI. However, to compare neural activity obtained with WOI to rs-fMRI, the experiments must be performed in mice. Fortunately, seminal work on the neural basis of fMRI and FC was performed in rodents and primates (*5, 6, 8, 41*), finding that the hemodynamic response is relatively conserved across species (*5, 6, 41–45*), and analogous functional networks can be observed in rodents, monkeys and humans (*46–48*) (review,(*49*)). We are therefore confident that the results of the WOI/rs-fMRI studies in mice are highly relevant to the rs-fMRI studies performed in humans. Although our experiments were performed under light isoflurane anesthesia while humans are typically awake, several lines of evidence suggest that the spatial structure that we measure represents a stable organizational backbone that is not strongly dependent on state (*7, 14, 49–53*). Future studies should certainly be performed in awake mice to confirm this. Both simultaneous WOI/rs-fMRI and rs-fMRI of awake mice are challenging experiments that will require further optimization prior to integration.

### Modularity vs. spatial similarity

Modularity has been used to describe the relative integration or segregation of the overall network structure of the brain. We had expected that modularity might provide a useful, high-level description of the changing patterns of cortical activity obtained with windowed approaches. However, modularity was fairly consistent across patterns that exhibited substantial spatial variability, and the time courses of modularity from different modalities showed little relationship to each other. The modularity analysis was originally designed for whole-brain images from humans, rather than the smaller matrices obtained from cortical images in mice, and may need further optimization to better characterize the changes that can be observed across time windows. We chose to focus on spatial similarity instead due to the ease of interpretability.

### Limitations

In addition to limitations discussed above, such as the use of anesthesia, a few other caveats should be noted. It is important to recall that all analysis other than the data quality demonstration in Figure 1 was performed using parcellated data rather than at the voxel/pixel level. Parcellation lends itself to network-based approaches such as FC and modularity, and is widely utilized in human rs-fMRI studies. However, parcellation is inherently insensitive to the fine details of spatial structure, which can be obscured by averaging all of time courses within the parcel. Future work would ideally compare FC from hemodynamics to FC from neural activity without parcellation to better characterize the similarity of the networks obtained.

While whole-brain rs-fMRI FC matrices were shown to provide context for the cortical FC, subcortical regions were not otherwise considered during analysis. Neural activity in these areas cannot be assessed with WOI, but may provide important context about overall brain state that helps to interpret cortical FC, a possibility that should be examined in the future.

Preprocessing methods for both rs-fMRI and WOI are still somewhat contentious. For rs-fMRI especially, the use of global signal regression continues to be debated (*54–57*). For this study, we took advantage of mouse anatomy to use signal from tissue adjacent to the brain as a regressor to remove widespread artifacts instead of the more controversial global signal regression (*58*). However, the mean signal was regressed from the WOI data to adjust for minor differences in overall light detection. Because GSR is known to influence anticorrelations (*55*), the small difference in preprocessing may account for the enhanced anticorrelation in WOI compared to rs-fMRI.

### Outlook

The development of simultaneous WOI and rs-fMRI enables direct comparison of rapidly-sampled fluorescence and HBT with the more slowly sampled hemodynamic BOLD signal, providing solid evidence that time-averaged rs-fMRI FC reflects the spatial structure of the underlying neural activity in the cortex. Moreover, the multimodal WOI/rs-fMRI approach will make it possible to optimize the sensitivity of rs-fMRI to neural activity using the “ground truth” fluorescence to assess the effects of alterations in rs-fMRI methods. This same approach can enable direct validation of other time-resolved analysis methods (CAPs, quasiperiodic patterns (*59–61*)), improving interpretation of rs-fMRI across species. Moreover, WOI is an extremely flexible modality that can be used to image activity in targeted subpopulations of neurons or even in glial cells such as astrocytes, providing unprecedented insight into the mechanisms that give rise to BOLD FC. These tools will be especially valuable for understanding alterations that occur with aging or neurodegeneration, where both neural activity and hemodynamics might be affected.

## Materials and Methods

### Experimental design

Simultaneous wide-field optical imaging and rs-fMRI (14 scans) were obtained from 9 mice expressing GCaMP6f in excitatory neurons and anesthetized with 1% isoflurane using a method reported previously (*16*). Image acquisition methods are summarized below for convenience. The simultaneously-acquired fluorescence, HBT, and rs-fMRI were parcellated and FC matrices were created for individual scans and for the whole group. For WOI, the analysis was repeated for three frequency bands (infraslow, slow, delta). The time-locked multimodal data were also segmented into shorter windows, with FC matrices for all modalities calculated for each window. Similarity of spatial structure was assessed across modalities and across conditions using spatial correlation of the FC matrices.

### Surgical preparation

All procedures were approved by the Emory Institutional Animal Care and Use Committee. Each mouse was anesthetized with isoflurane (2-3%) and a headpiece with an open center ∼1 cm in diameter was surgically attached with dental cement. The skull within the opening was thinned and a round glass coverslip was attached using optical adhesive to create a crystal-clear and long-lasting optical window. After surgery, each mouse was allowed to recover for at least one week prior to imaging.

### Imaging

All WOI and rs-fMRI was performed in the bore of a 9.4 T/20 cm preclinical MRI system (Bruker, Billerica, MA). Mice were anesthetized with 2-3% isoflurane for setup and maintained at approximately 1% during imaging. A transmit/receive surface coil, shaped to fit the mouse brain, was positioned around the implanted headpiece, which is held in place by a fitted bar. The WOI system includes LED light delivered to the brain by optical fibers and a conventional camera-based detection approach that utilizes lens and mirrors to direct the light to the cameras situated in the control room outside of the scanner. Light was delivered at 466 nm for excitation of the fluorophor and at 525 nm for reflectance imaging of total hemoglobin (HBT), which is proportional to cerebral blood volume. The emitted and reflected light (both 525 nm) were detected in interleaved fashion at a rate of 50 Hz for each signal using a Kuros camera situated in the control room.

After positioning and shimming on the region of interest, anatomical (FLASH, 1mm slices, 130 micron x 130 micron spatial resolution) and time-of-flight MR images (FLASH, 14 coronal slices, 200 micron thickness, 69 x 69 micron in plane resolution) were obtained, then rs-fMRI (GE-EPI; TR 1 s, TE 15 ms, 400 micron isotropic resolution) and WOI (fluorescence and reflectance, 50 Hz each, ∼1 cm^2^ field of view, 100 x 100 matrix) were acquired for 20 minutes. Temperature, heart rate, and respiration were maintained within normal physiological ranges. Physiological parameters and TTL pulses for each imaging frame from the MRI and camera system were recorded in Matlab. Scans from multiple days for three mice (2 days for two mice, 3 days for one mouse) are included in the dataset.

### Pre-processing

Data for each modality were preprocessed separately. Preprocessing for rs-fMRI included motion correction, regression of tissue signal from outside the brain (to remove large-scale artifacts while minimizing loss of brain activity (*58*)), smoothing, and bandpass filtering (0.01-0.1 Hz). The WOI images were smoothed and global signal regression was used to minimize large-scale changes. Three separate bandpass filters (infraslow, 0.01-0.1 Hz; slow, 0.1 – 1 Hz; delta, 1-4 Hz) were applied to WOI data to enable frequency-specific analysis. WOI of fluorescence was corrected for hemodynamic contamination using regression of the HBT images (both z-scored). WOI and rs-fMRI data were aligned to the Allen mouse brain atlas (*62*). The cranial window covered most but not all of the areas of the Allen cortical atlas; areas without coverage were not included in the analysis. The areas covered varied slightly across mice; for group analysis, only areas present for all mice were considered. For each parcel within the cortical atlas, the rs-fMRI signal was summed from the entire 3D volume of that parcel rather than the surface alone (as for WOI). For the time-varying analysis, data were manually aligned in time using the recorded time-locked TTL signals. Any segments containing artifacts or motion in the MRI scan (none were evident in the WOI data) were discarded prior to further analysis.

### Analysis of FC

Functional connectivity was obtained from unparcellated data for each modality using ICA (GIFT toolbox, 20 components) to demonstrate data quality. After parcellation, a functional connectivity (FC) matrix was then calculated for each modality as the pairwise correlation between the average signal time course for every parcel from the cortical surface. FC matrices also calculated for all parcels in the 3D Allen brain atlas for the rs-fMRI signal as context for the cortical FC matrices. These FC matrices were calculated at the group level and at the individual scan level. The same analysis was performed for WOI data filtered into slow and delta frequency bands to determine whether spatial structure was altered across frequencies. No frequencies above 4 Hz were examined. As shown in **Supplemental Figure 1**, preliminary analysis from individual scans indicated that FC from fluorescence lost most of its spatial structure at higher frequencies, which is to expected from the finite time constant of the calcium indicator.

Similarity of FC matrices was measured using spatial correlation, a metric that is sensitive to the spatial pattern of the signal but not to the relative amplitude of the pattern, which we anticipated may vary across modalities. Using this approach, similarity of FC matrices was characterized across modalities in the same mice at the group and individual level. Similarity of FC matrices was also examined for the same modalities obtained from the same mice on different days (up to 3 scans per mouse).

### Analysis of time-resolved FC

For time-resolved analysis, time series from all modalities were divided into non-overlapping windows 15 seconds, 30 seconds, or 60 seconds in length. For delta-band WOI alone, a window of 1 second was also used. FC matrices were calculated for each window, and similarity was assessed using spatial correlation within and across modalities.

### Modularity

The organization of brain-wide intrinsic activity has sometimes been characterized in terms of relative integration or segregation. To obtain some insight into this property of the brain’s network structure, modularity of group average and individual FC matrices was assessed iteratively using the Louvain algorithm in the Brain Connectivity Toolbox (*63*), with negative correlation treated symmetrically and gamma = 1.

### Statistical analysis

Comparisons of modularity and spatial similarity were performed using a two-sided t-test, with p<0.05 considered significant. Correction for multiple comparisons followed the Bonferroni method, with n=3 for comparisons across modalities (fluorescence to HBT, HBT to rs-fMRI, rs-fMRI to fluorescence).

## Funding

Emory University Center for Systems Imaging Core

NIH 1S10OD028503

NIH 1R21MH140191

NIH 1R01NS078095

NIH R21NS122013

## Author contributions

Conceptualization: W-JP, SK

Methodology: W-JP, SK

Investigation: W-JP, LD, LM-B, SK

Visualization: W-JP, SK

Supervision: SK

Writing—original draft: SK, W-JP

Writing—review & editing: W-JP, SK, LD, LM-B

## Data and materials availability

All data are available in the main text or the supplementary materials. The data and code will be publicly available at Zenodo with the identifier [DOI will be added upon reviewer-access/acceptance].

---

**Supplemental Figure 1.**
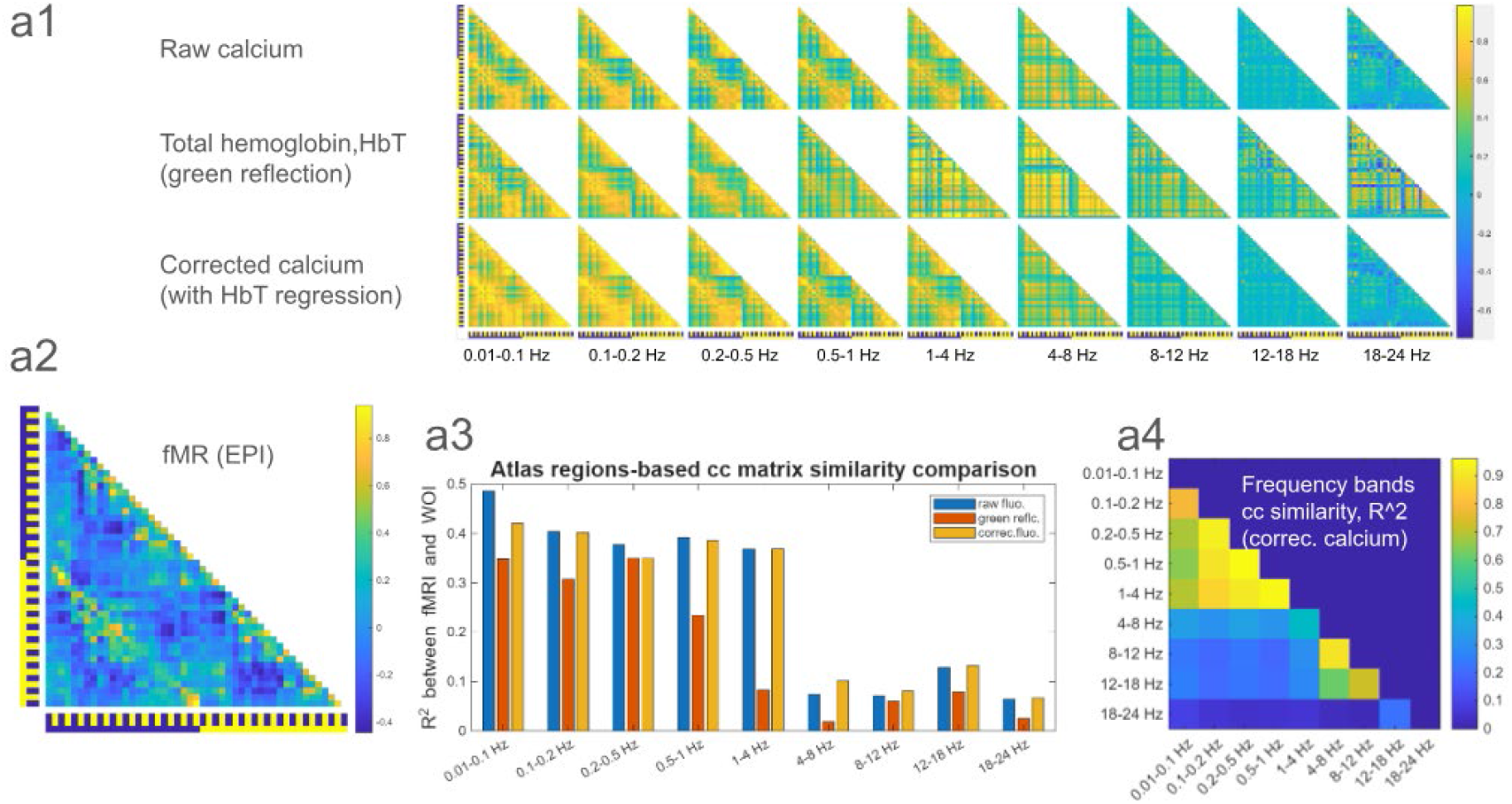
Preliminary analysis of unparcellated data from one mouse. FC matrices were created for raw fluorescence, HBT, and corrected fluorescence images filtered into a variety of frequency bands. At frequencies above 4 Hz, little spatial structure is observed for any modality.

**Supplemental Figure 2.**
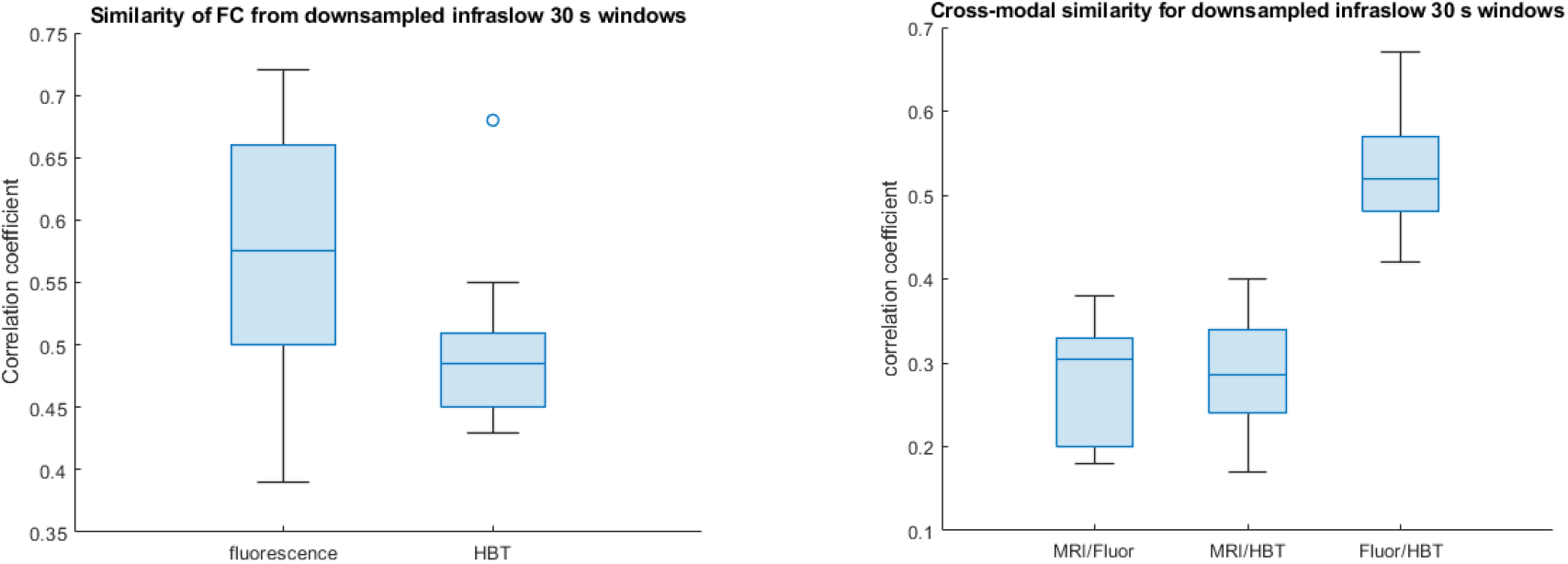
A) Similarity of FC in 30 second windows from infraslow fluorescence and HBT after downsampling to the temporal resolution of the rs-fMRI signal (1 Hz). B) Similarity of FC matrices from the same window across modalities, using downsampled WOI data from the infraslow band with a 30 second window. No significant differences between the similarity of FC matrices created from the downsampled data and the similarity of the FC matrices created from the original data (Figure 7) were observed, suggesting that the lower sampling rate alone is not sufficient to account for the differences observed for rs-fMRI.

**Supplemental Figure 3.**
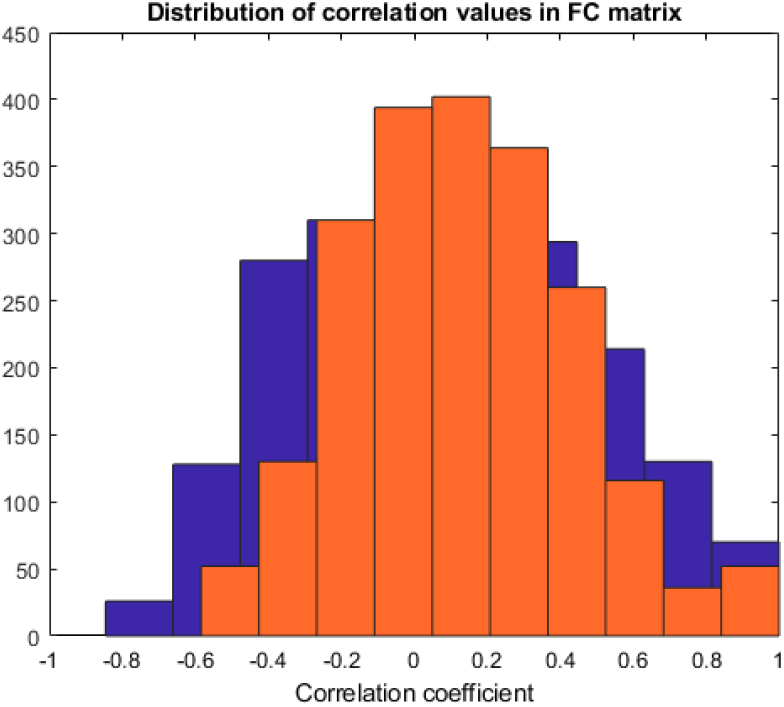
Distribution of correlation coefficients from a FC matrix showing high similarity to fluorescence FC and structured intra- and inter-hemispheric connectivity (blue) and from a FC matrix with low similarity to fluorescence FC and loss of structured connectivity (orange). The histogram is narrower and centered near zero for the FC matrix with little structure, suggesting that it might be possible to infer windows with poor similarity based upon the shape of their histogram or other related features.

**Supplemental Figure 4.**
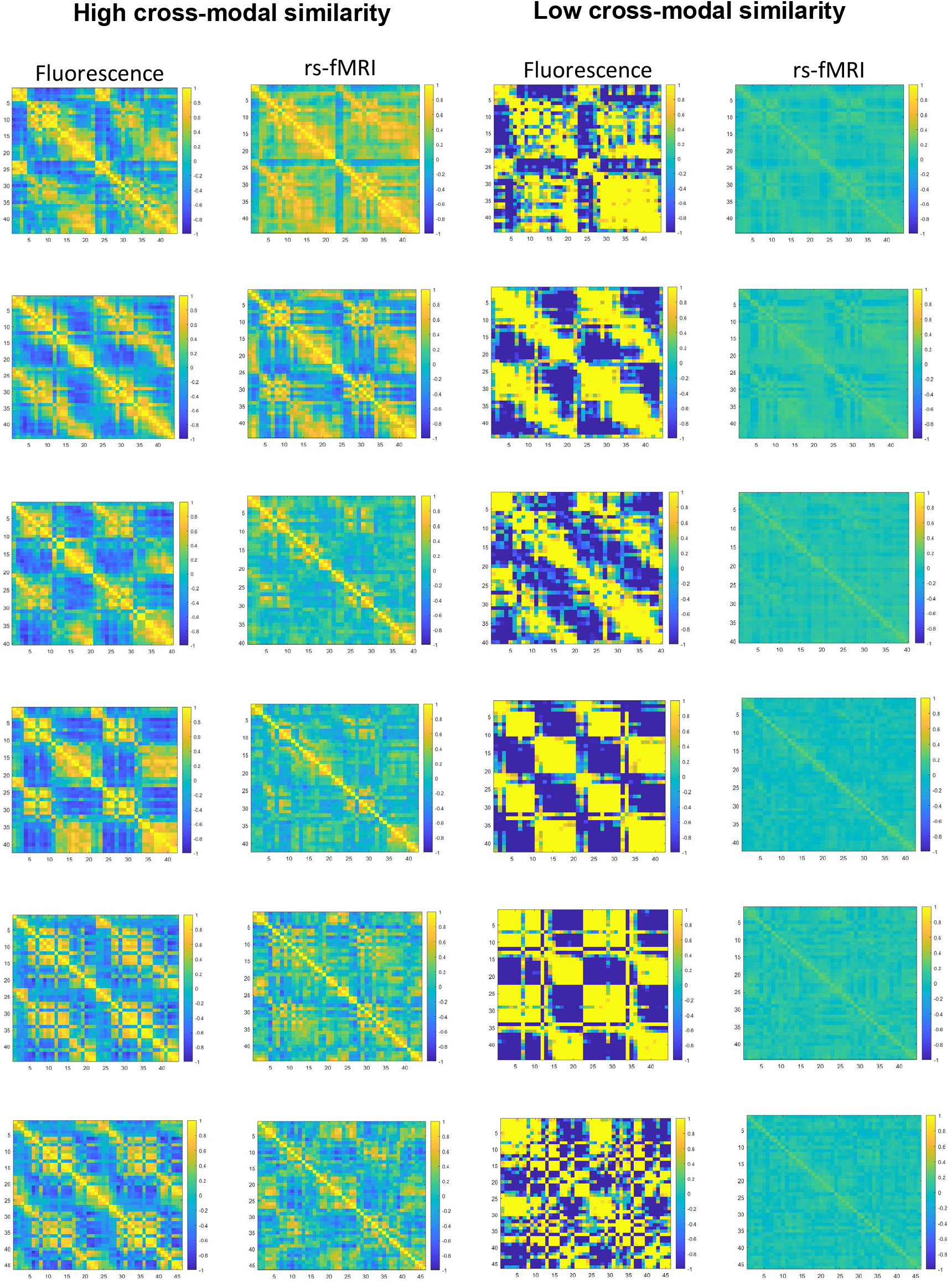
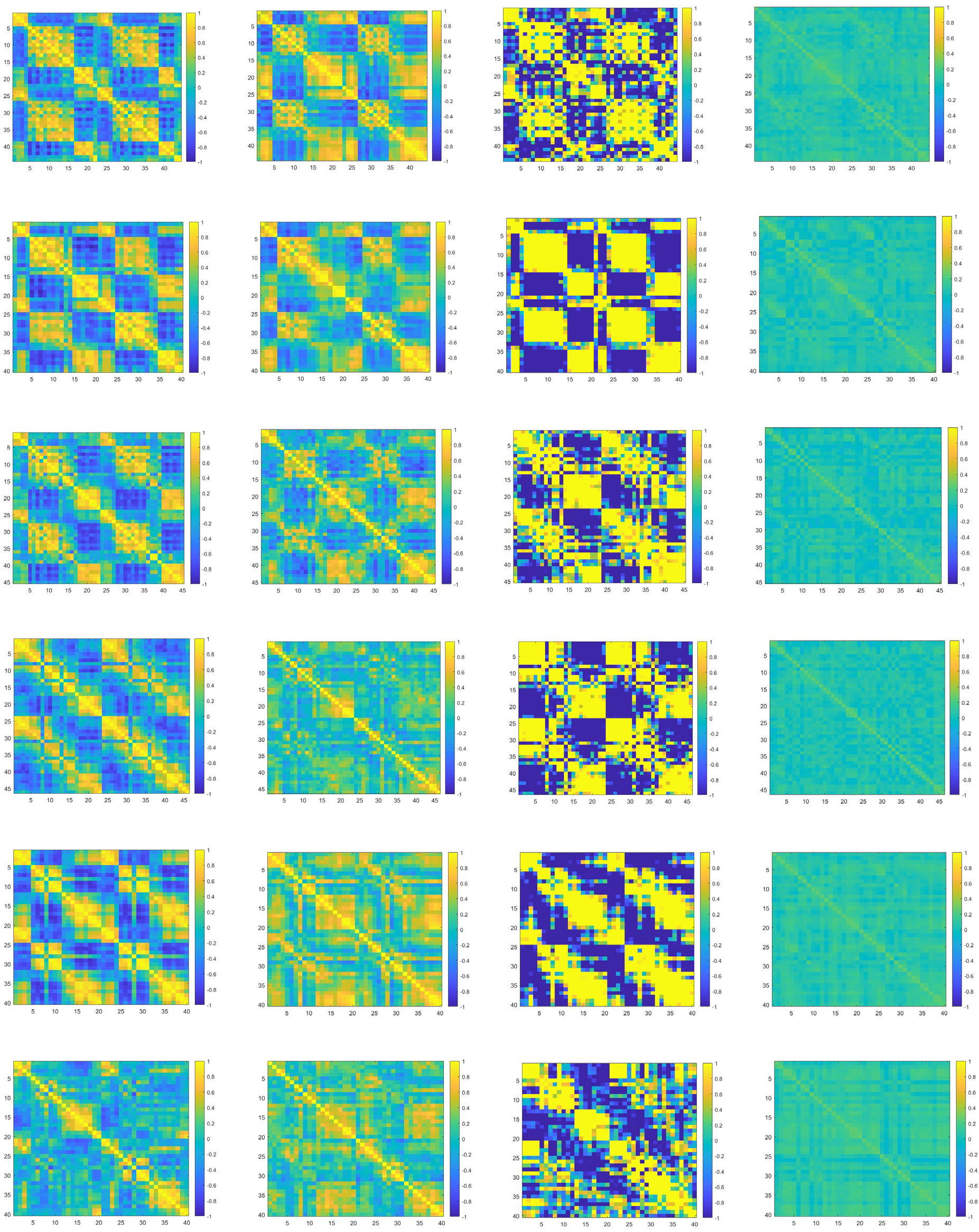
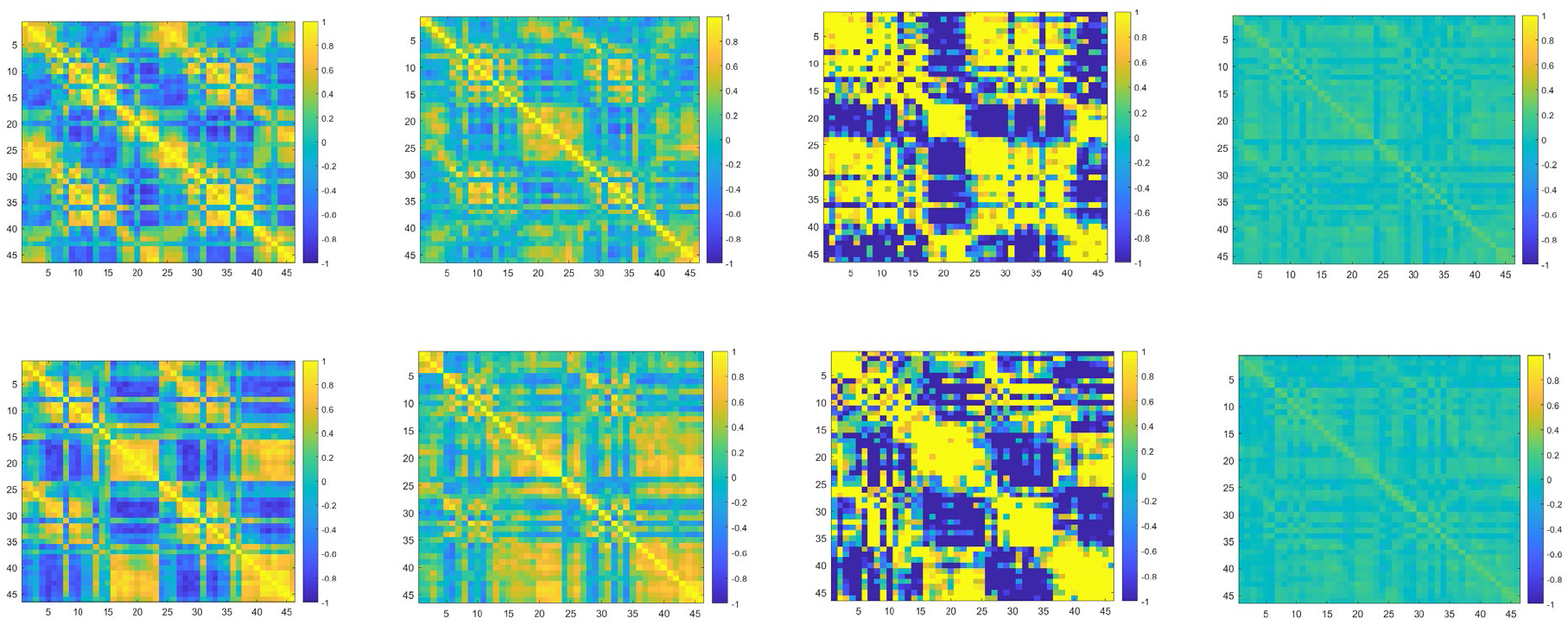
FC matrices created from windows with the highest and lowest similarity between rs-fMRI and fluorescence for each scan (one per row).

## References

1. B. Biswal, F. Z. Yetkin, V. M. Haughton, J. S. Hyde, Functional connectivity in the motor cortex of resting human brain using echo-planar MRI. Magn Reson Med 34, 537–541 (1995).

2. R. M. M. Hutchison, T. Womelsdorf, E. A. E. A. Allen, P. A. P. A. Bandettini, V. D. V. D. Calhoun, M. Corbetta, S. Della Penna, J. H. J. H. Duyn, G. H. G. H. Glover, J. Gonzalez-Castillo, D. A. D. A. Handwerker, S. Keilholz, V. Kiviniemi, D. A. D. A. Leopold, F. de Pasquale, O. Sporns, M. Walter, C. Chang, Dynamic functional connectivity: promise, issues, and interpretations. NeuroImage 80, 360–378 (2013).

3. M. G. Preti, T. A. Bolton, D. Van De Ville, The dynamic functional connectome: State-of-the-art and perspectives. NeuroImage 160, 41–54 (2017).

4. S. Keilholz, C. Caballero-Gaudes, P. Bandettini, G. Deco, V. Calhoun, Time-Resolved Resting-State Functional Magnetic Resonance Imaging Analysis: Current Status, Challenges, and New Directions. Brain Connectivity 7, 465–481 (2017).

5. N. K. Logothetis, J. Pauls, M. Augath, T. Trinath, A. Oeltermann, Neurophysiological investigation of the basis of the fMRI signal. Nature 412, 150–157 (2001).

6. A. Shmuel, D. A. Leopold, Neuronal correlates of spontaneous fluctuations in fMRI signals in monkey visual cortex: Implications for functional connectivity at rest. Hum Brain Mapp 29, 751–761 (2008).

7. W.-J. J. W.-J. Pan, G. J. G. J. Thompson, M. E. M. E. Magnuson, D. Jaeger, S. Keilholz, Infraslow LFP correlates to resting-state fMRI BOLD signals. NeuroImage 74, 288–297 (2013).

8. W.-J. Pan, G. Thompson, M. Magnuson, W. Majeed, D. Jaeger, S. Keilholz, Broadband Local Field Potentials Correlate with Spontaneous Fluctuations in Functional Magnetic Resonance Imaging Signals in the Rat Somatosensory Cortex Under Isoflurane Anesthesia. Brain Connectivity 1, 119–131 (2011).

9. H. Lu, Y. Zuo, H. Gu, J. A. Waltz, W. Zhan, C. A. Scholl, W. Rea, Y. Yang, E. A. Stein, Synchronized delta oscillations correlate with the resting-state functional MRI signal. Proc Natl Acad Sci U S A 104, 18265–18269 (2007).

10. Y. Ma, M. A. Shaik, M. G. Kozberg, S. H. Kim, J. P. Portes, D. Timerman, E. M. C. C. Hillman, Resting-state hemodynamics are spatiotemporally coupled to synchronized and symmetric neural activity in excitatory neurons. Proceedings of the National Academy of Sciences of the United States of America 113, 201525369 (2016).

11. P. Pais-Roldán, C. Mateo, W.-J. Pan, B. Acland, D. Kleinfeld, L. H. Snyder, X. Yu, S. Keilholz, Contribution of animal models toward understanding resting state functional connectivity. NeuroImage 245, 118630 (2021).

12. G. J. Thompson, W.-J. Pan, S. D. Keilholz, Different dynamic resting state fMRI patterns are linked to different frequencies of neural activity. Journal of neurophysiology 114, 114–24 (2015).

13. H. Uhlirova, K. Kılıç, P. Tian, M. Thunemann, M. Desjardins, P. A. Saisan, S. Sakadžić, T. V. Ness, C. Mateo, Q. Cheng, K. L. Weldy, F. Razoux, M. Vandenberghe, J. A. Cremonesi, C. G. Ferri, K. Nizar, V. B. Sridhar, T. C. Steed, M. Abashin, Y. Fainman, E. Masliah, S. Djurovic, O. A. Andreassen, G. A. Silva, D. A. Boas, D. Kleinfeld, R. B. Buxton, G. T. Einevoll, A. M. Dale, A. Devor, Cell type specificity of neurovascular coupling in cerebral cortex. Elife 5, e14315 (2016).

14. P. W. Wright, L. M. Brier, A. Q. Bauer, G. A. Baxter, A. W. Kraft, M. D. Reisman, A. R. Bice, A. Z. Snyder, J. M. Lee, J. P. Culver, Functional connectivity structure of cortical calcium dynamics in anesthetized and awake mice. PLoS ONE 12 (2017).

15. E. M. R. Lake, X. Ge, X. Shen, P. Herman, F. Hyder, J. A. Cardin, M. J. Higley, D. Scheinost, X. Papademetris, M. C. Crair, R. T. Constable, Simultaneous cortex-wide fluorescence Ca2+ imaging and whole-brain fMRI. Nature methods 17, 1262–1271 (2020).

16. W.-J. Pan, L. Daley, H. Watters, L. Meyer-Baese, K. Gopinath, D. Jaeger, S. Keilholz, An integrated platform for simultaneous wide-field voltage/calcium imaging and fMRI (EPI &; ZTE) reveals neuronal infraslow dynamics underlying functional connectivity. bioRxiv [Preprint] (2026). 10.64898/2026.01.26.701889.

17. A. W. Chan, M. H. Mohajerani, J. M. LeDue, Y. T. Wang, T. H. Murphy, Mesoscale infraslow spontaneous membrane potential fluctuations recapitulate high-frequency activity cortical motifs. Nature communications 6, 7738 (2015).

18. Y. Ma, M. A. Shaik, S. H. Kim, M. G. Kozberg, D. N. Thibodeaux, H. T. Zhao, H. Yu, E. M. C. C. Hillman, Wide-Field Optical Mapping of Neural Activity and Brain Haemodynamics: Considerations and Novel Approaches (Royal Society of London, 2016; http://rstb.royalsocietypublishing.org/lookup/doi/10.1098/rstb.2015.0360 )vol. 371.

19. S. Keilholz, M. E. Magnuson, W.-J.-.J. Pan, M. Willis, G. Thompson, Dynamic Properties of Functional Connectivity in the Rodent. Brain Connect, 121029090330004 (2012).

20. S. Shakil, C. H. C.-H. Lee, S. D. S. D. Keilholz, Evaluation of sliding window correlation performance for characterizing dynamic functional connectivity and brain states. NeuroImage 133, 111–128 (2016).

21. R. Hindriks, M. H. Adhikari, Y. Murayama, M. Ganzetti, D. Mantini, N. K. Logothetis, G. Deco, Can sliding-window correlations reveal dynamic functional connectivity in resting-state fMRI? NeuroImage 127, 242–256 (2015).

22. A. M. Mishra, D. J. Ellens, U. Schridde, J. E. Motelow, M. J. Purcaro, M. N. DeSalvo, M. Enev, B. G. Sanganahalli, F. Hyder, H. Blumenfeld, Where fMRI and electrophysiology agree to disagree: corticothalamic and striatal activity patterns in the WAG/Rij rat. J Neurosci 31, 15053–15064 (2011).

23. B.-X. B.-X. Huo, J. B. Smith, P. J. Drew, Neurovascular coupling and decoupling in the cortex during voluntary locomotion. The Journal of neuroscience : the official journal of the Society for Neuroscience 34, 10975–81 (2014).

24. I. M. Devonshire, N. G. Papadakis, M. Port, J. Berwick, A. J. Kennerley, J. E. W. Mayhew, P. G. Overton, Neurovascular coupling is brain region-dependent. NeuroImage 59, 1997–2006 (2012).

25. R. M. Birn, J. B. Diamond, M. A. Smith, P. A. Bandettini, Separating respiratory-variation-related fluctuations from neuronal-activity-related fluctuations in fMRI. NeuroImage 31, 1536–1548 (2006).

26. J. D. Power, K. A. Barnes, A. Z. Snyder, B. L. Schlaggar, S. E. Petersen, K. A. Barnes, S. E. Petersen, J. D. Power, B. L. Schlaggar, Spurious but systematic correlations in functional connectivity MRI networks arise from subject motion. NeuroImage 59, 2142–2154 (2012).

27. J. D. Power, M. Plitt, T. O. Laumann, A. Martin, Sources and implications of whole-brain fMRI signals in humans. NeuroImage 146, 609–625 (2017).

28. T. T. Liu, Noise contributions to the fMRI signal: An overview. NeuroImage 143, 141–151 (2016).

29. S. Moeller, P. K. Pisharady, S. Ramanna, C. Lenglet, X. Wu, L. Dowdle, E. Yacoub, K. Uğurbil, M. Akçakaya, NOise reduction with DIstribution Corrected (NORDIC) PCA in dMRI with complex-valued parameter-free locally low-rank processing. NeuroImage 226, 2020.08.25.267062 (2021).

30. R. W. Chan, G. Hamilton-Fletcher, B. J. Edelman, M. A. Faiq, T. A. Sajitha, S. Moeller, K. C. Chan, NOise Reduction with DIstribution Corrected (NORDIC) principal component analysis improves brain activity detection across rodent and human functional MRI contexts. Imaging Neuroscience, doi: 10.1162/imag_a_00325 (2024).

31. C. Chang, G. H. H. Glover, Time-frequency dynamics of resting-state brain connectivity measured with fMRI. NeuroImage 50, 81–98 (2010).

32. E. A. Allen, E. Damaraju, S. M. Plis, E. B. Erhardt, T. Eichele, V. D. Calhoun, Tracking whole-brain connectivity dynamics in the resting state. Cereb Cortex 24, 663–676 (2014).

33. N. Leonardi, D. Van De Ville, On spurious and real fluctuations of dynamic functional connectivity during rest. NeuroImage 104, 430–436 (2015).

34. G. J. G. J. Thompson, M. D. M. D. Merritt, W.-J. W. J. Pan, M. E. M. E. Magnuson, J. K. J. K. Grooms, D. Jaeger, S. D. S. D. Keilholz, Neural correlates of time-varying functional connectivity in the rat. NeuroImage 83, 826–836 (2013).

35. J. Gonzalez-Castillo, C. W. Hoy, D. A. Handwerker, M. E. Robinson, L. C. Buchanan, Z. S. Saad, P. A. Bandettini, Tracking ongoing cognition in individuals using brief, whole-brain functional connectivity patterns. Proceedings of the National Academy of Sciences of the United States of America, 1501242112– (2015).

36. X. Liu, J. H. H. Duyn, Time-varying functional network information extracted from brief instances of spontaneous brain activity. Proc Natl Acad Sci U S A 110, 4392–4397 (2013).

37. N. Petridou, C. C. Gaudes, I. L. Dryden, S. T. Francis, P. A. Gowland, Periods of rest in fMRI contain individual spontaneous events which are related to slowly fluctuating spontaneous activity. Human Brain Mapping 34, 1319–1329 (2013).

38. P. Figueiredo, I. Esteves, A. Fouto, A. Ruiz-Tagle, G. Caetano, C. Caballero-Gaudes, Electrophysiological correlates of BOLD events with high cofluctuation amplitude in the resting human brain.

39. G. J. Thompson, W. J. Pan, M. E. Magnuson, D. Jaeger, S. D. Keilholz, Quasi-periodic patterns (QPP): Large-scale dynamics in resting state fMRI that correlate with local infraslow electrical activity. NeuroImage 84, 1018–1031 (2014).

40. G. J. Thompson, W.-J. Pan, J. C. W. Billings, J. K. K. Grooms, S. Shakil, D. Jaeger, S. D. Keilholz, Phase-amplitude coupling and infraslow (<1 Hz) frequencies in the rat brain: relationship to resting state fMRI. Front Integr Neurosci 8, 41 (2014).

41. G. Brinker, C. Bock, E. Busch, H. Krep, K. A. Hossmann, M. Hoehn-Berlage, Simultaneous recording of evoked potentials and T2*-weighted MR images during somatosensory stimulation of rat. Magn Reson Med 41, 469–473 (1999).

42. Y. Nir, R. Mukamel, I. Dinstein, E. Privman, M. Harel, L. Fisch, H. Gelbard-Sagiv, S. Kipervasser, F. Andelman, M. Y. Neufeld, U. Kramer, A. Arieli, I. Fried, R. Malach, Interhemispheric correlations of slow spontaneous neuronal fluctuations revealed in human sensory cortex. Nat Neurosci 11, 1100–1108 (2008).

43. B. J. J. He, A. Z. Z. Snyder, J. M. M. Zempel, M. D. D. Smyth, M. E. E. Raichle, Electrophysiological correlates of the brain’s intrinsic large-scale functional architecture. Proc Natl Acad Sci U S A 105, 16039–16044 (2008).

44. J. A. de Zwart, A. C. Silva, P. van Gelderen, P. Kellman, M. Fukunaga, R. Chu, A. P. Koretsky, J. A. Frank, J. H. Duyn, Temporal dynamics of the BOLD fMRI impulse response. Neuroimage 24, 667–677 (2005).

45. N. Hewson-Stoate, M. Jones, J. Martindale, J. Berwick, J. Mayhew, Further nonlinearities in neurovascular coupling in rodent barrel cortex. Neuroimage 24, 565–574 (2005).

46. H. Lu, Q. Zou, H. Gu, M. E. Raichle, E. A. Stein, Y. Yang, Rat brains also have a default mode network. Proc Natl Acad Sci U S A 109, 3979–3984 (2012).

47. M. D. D. Fox, A. Z. Z. Snyder, J. L. L. Vincent, M. Corbetta, D. C. C. Van Essen, M. E. E. Raichle, The human brain is intrinsically organized into dynamic, anticorrelated functional networks. Proc Natl Acad Sci U S A 102, 9673–9678 (2005).

48. D. Mantini, A. Gerits, K. Nelissen, J.-B. Durand, O. Joly, L. Simone, H. Sawamura, C. Wardak, G. A. Orban, R. L. Buckner, W. Vanduffel, Default mode of brain function in monkeys. J Neurosci 31, 12954–12962 (2011).

49. N. Xu, T. J. LaGrow, N. Anumba, A. Lee, X. Zhang, B. Yousefi, Y. Bassil, G. P. Clavijo, V. Khalilzad Sharghi, E. Maltbie, L. Meyer-Baese, M. Nezafati, W.-J. Pan, S. Keilholz, Functional Connectivity of the Brain Across Rodents and Humans. Frontiers in Neuroscience 16 (2022).

50. D. Gutierrez-Barragan, M. A. Basson, S. Panzeri, A. Gozzi, Infraslow State Fluctuations Govern Spontaneous fMRI Network Dynamics. Current biology : CB 29, 2295–2306.e5 (2019).

51. D. N. Guilfoyle, S. V. Gerum, J. L. Sanchez, A. Balla, H. Sershen, D. C. Javitt, M. J. Hoptman, Functional connectivity fMRI in mouse brain at 7T using isoflurane. Journal of neuroscience methods 214, 144–8 (2013).

52. K. Masamoto, T. Kim, M. Fukuda, P. Wang, S. G. Kim, Relationship between neural, vascular, and BOLD signals in isoflurane-anesthetized rat somatosensory cortex. Cereb Cortex 17, 942–950 (2006).

53. A. Gozzi, Evolutionarily conserved fMRI network dynamics in the mouse, macaque, and human brain. Nature Communications 15 (2024).

54. J. C. W. Billings, S. D. Keilholz, The Not-So-Global BOLD Signal. Brain Connectivity, brain.2017.0517 (2018).

55. K. Murphy, R. M. Birn, D. A. Handwerker, T. B. Jones, P. A. Bandettini, The impact of global signal regression on resting state correlations: Are anti-correlated networks introduced? NeuroImage 44, 893–905 (2009).

56. K. Murphy, M. D. Fox, Towards a consensus regarding global signal regression for resting state functional connectivity MRI. NeuroImage 154, 169–173 (2017).

57. Z. S. Saad, S. J. Gotts, K. Murphy, G. Chen, H. J. Jo, A. Martin, R. W. Cox, Trouble at rest: how correlation patterns and group differences become distorted after global signal regression. Brain connectivity 2, 25–32 (2012).

58. N. Anumba, E. Maltbie, W.-J. Pan, T. J. LaGrow, N. Xu, S. Keilholz, Spatial and Spectral Components of the BOLD Global Signal in Rat Resting-State Functional MRI. Magnetic Resonance in Medicine 90, 2486–2499 (2023).

59. W. Majeed, M. Magnuson, W. Hasenkamp, H. Schwarb, E. H. E. H. H. Schumacher, L. Barsalou, S. D. D. S. D. Keilholz, Spatiotemporal dynamics of low frequency BOLD fluctuations in rats and humans. NeuroImage 54, 1140–1150 (2011).

60. B. Yousefi, S. Keilholz, Propagating patterns of intrinsic activity along macroscale gradients coordinate functional connections across the whole brain. NeuroImage 231, 117827 (2021).

61. T. Bolt, J. S. Nomi, D. Bzdok, C. Chang, B. T. T. Yeo, L. Q. Uddin, S. D. Keilholz, A Parsimonious Description of Global Functional Brain Organization in Three Spatiotemporal Patterns. bioRxiv, 2021.06.20.448984 (2022).

62. S. W. Oh, J. A. Harris, L. Ng, B. Winslow, N. Cain, S. Mihalas, Q. Wang, C. Lau, L. Kuan, A. M. Henry, M. T. Mortrud, B. Ouellette, T. N. Nguyen, S. A. Sorensen, C. R. Slaughterbeck, W. Wakeman, Y. Li, D. Feng, A. Ho, E. Nicholas, K. E. Hirokawa, P. Bohn, K. M. Joines, H. Peng, M. J. Hawrylycz, J. W. Phillips, J. G. Hohmann, P. Wohnoutka, C. R. Gerfen, C. Koch, A. Bernard, C. Dang, A. R. Jones, H. Zeng, A mesoscale connectome of the mouse brain. Nature 508, 207–214 (2014).

63. M. Rubinov, O. Sporns, Complex network measures of brain connectivity: uses and interpretations. NeuroImage 52, 1059–1069 (2010).

